# Integrating citizen science and state agency data expands environmental coverage of occurrence data in invasive species distribution models

**DOI:** 10.64898/2026.04.20.719594

**Authors:** A. Horn, V. Lozano, T. Kleinebecker, Y.P. Klinger

**Affiliations:** Division of Landscape Ecology and Landscape Planning, Justus Liebig University, Heinrich-Buff-Ring 26-32, 35392 Gießen, Germany; Department of Agricultural Sciences, University of Sassari, Viale Italia 39/A, 07100 Sassari, Italy

**Author notes:** Corresponding author:* Yves P. Klinger.

**Keywords:** invasive plants, species distribution models, niche overlap, environmental space, habitat suitability, area of model applicability

## Abstract

Species distribution models (SDMs) are widely used to support risk assessment for invasive non-native plant species (invasive plants), but their performance is constrained by the coverage of occurrence data. Combining occurrences from citizen science (CS) platforms with data from state agency (StAg) monitoring may provide advantages, as these datasets follow different sampling strategies, yet they are rarely integrated. Here, we systematically compare how CS, StAg, and combined data influence the inferred environmental spaces, predictive discrimination, and spatial applicability of SDMs for three widespread invasive plants (*A. altissima*, *H. mantegazzianum*, *I. glandulifera*) in central Germany. We quantified overlap of sampled environmental space between datasets using PCA and Schoener’s D and applied a hierarchical SDM utilizing boosted regression trees. We then assessed the Area of Applicability to identify environmental conditions outside of the training data for either model, which could represent priorities for additional monitoring.

CS data were biased toward lower-elevation, urbanized environments, whereas StAg data captured higher-elevation, remote habitats, particularly along watercourses. Overlap was lowest for *A. altissima*, which is spreading, intermediate for *H. mantegazzianum*, associated with linear habitats, and highest for *I. glandulifera*, a widespread species, which may reflect differences in invasion stage and habitat associations as well as in sampling. Combined models had slightly higher mean AUC (0.85 ± 0.07) and TSS (0.58 ± 0.15), but overall discrimination did not differ significantly among data sources, and the best-performing source differed among species. Applicability (≥59%) was high across data sources and species, with combined models showing weaker increases in environmental dissimilarity along environmental gradients. Thus, CS and StAg records sampled distinct parts of the environmental space, a difference reflected in suitability patterns and extrapolation but not detected by AUC and TSS. Integrating both sources broadened the environmental representation of occurrence records without removing their individual biases, which is relevant for invasive plants risk assessments.

**Highlights:**

- Citizen science and state agency data capture distinct environmental spaces.
- Differences between data sources are related to sampling processes.
- Mean AUC/TSS was slightly higher for combined data, but not significantly so.
- State agency habitat mapping complements citizen science in invasive plant models.
- Underrepresented remote, high-elevation areas may be priorities for monitoring.

## 1. Introduction

The number of invasive non-native plant species continues to increase globally, while many established invasive plants expand their ranges (Roy et al., 2024; Seebens et al., 2017). This poses substantial challenges for biodiversity, human health, and the economy (Maxwell et al., 2016; Pyšek et al., 2020). In response, international policy frameworks like Target 6 of the Kunming-Montreal Biodiversity Framework, EU Regulation 1143/2014 on invasive alien species or national plans such as Germany’s action plan on the pathways of invasive species (Mayer et al., 2023) emphasize prevention, early detection and prioritization of management for invasive plants. To support these objectives, a range of frameworks and tools have been developed, including horizon scanning to anticipate emerging invasions (Roy et al., 2014), and novel technologies designed to identify invasive species at early stages of establishment (Martinez et al., 2020; Sepulveda et al., 2023). Among these tools, species distribution models (SDMs) are a well-established and widely used approach. Through spatial projections, they can support invasive plant risk assessment and management in a variety of ways, including assessing estimates of the spread of invasive plants under current and future environments (Anibaba et al., 2022; Barbet-Massin et al., 2018) and supporting prioritisation of invasive plants and sites at high invasion risk (Lozano et al., 2024; McGeoch et al., 2016). Despite their usefulness, SDMs are fundamentally constrained by the amount, quality, and coverage of species occurrence data (Moudrý et al., 2024; Tessarolo et al., 2021). The area of applicability (AOA) addresses this limitation by delineating where a model is applicable, i.e. where the predictor values are similar enough to the training data, based on the dissimilarity of predictor values to the training data (Meyer and Pebesma, 2021). While the AOA is increasingly used to communicate where predictions are applicable, it has rarely been used to compare models built on different data sources, or to link extrapolation to specific environmental gradients and data sources.

The limitation of SDMs is particularly pronounced for invasive species, which are often not in equilibrium with their invaded environments due to ongoing spreads, time lags, and dispersal limits (Gallien et al., 2012; Václavík and Meentemeyer, 2012). Consequently, SDMs for invasive plants are sensitive to how, where, and by whom occurrence data is collected. Occurrence data used to calibrate SDMs is predominantly obtained from biodiversity databases which include substantial contributions from citizen science (CS). Platforms such as the Global Biodiversity Information Facility (GBIF) are among the most widely used sources to obtain occurrence data used in SDMs (Feldman et al., 2021). These databases provide broad species coverage and large volumes of occurrence data at low cost, making them an attractive for scientists. Previous studies demonstrate that models derived from CS occurrence data can produce ecologically meaningful results, provided the inherent limitations of those datasets are considered appropriately (Johnston et al., 2020; Klinger et al., 2023; Matutini et al., 2021). Nonetheless, CS data may fail to capture the full extent of invasive plant niches due to misidentification and spatial or temporal bias (Cox et al., 2012; Feldman et al., 2021; Geldmann et al., 2016). Previous research has shown that integrating CS and expert-based datasets can improve on these biases and better SDM performance, as it expands spatial and environmental coverage and increases the accuracy of model results (Crall et al., 2015; Dimson et al., 2023; Robinson et al., 2020). Such studies have shown that professional and CS data can have inverse spatial biases. However, how these differences translate into the sampled environmental space, and into model applicability and extrapolation, has to our knowledge not been quantified.

In Europe, for invasive non-native species of Union Concern (EU Regulation 1143/2014), there is a legal mandate for monitoring and reporting by EU member states, resulting in large occurrence datasets collected by state agencies (StAg). How this data is collected differs between regions and member states and StAg data itself may result from activities such as habitat mapping, leading to recording biases. Initiatives such as the European Alien Species Information Network (EASIN, 2026) aim to centralize occurrence information on invasive plants, including data generated by StAg monitoring programs. Yet, despite these monitoring obligations, StAg data remains scarcely available and thus underutilized in ecological modelling. Even when accessible, StAg datasets cannot be considered a neutral baseline, as they are subject to biases reflecting the national or regional monitoring strategies. Currently, the large-scale aggregation and incomplete representation of national monitoring data strongly constrain their usability, leaving many StAg occurrence datasets difficult to access and rarely integrated into ecological models. Nevertheless, combining StAg and CS data in SDMs could leverage the strengths of both sources while mitigating individual biases, thus enhancing spatial coverage and improving the representation of the environmental space occupied by the species.

Despite monitoring obligations across EU member states, the contribution of StAg data to invasive plants environmental space characterization and the benefits of its integration into CS occurrence data for SDM predictions has not been quantified. We address this gap with a framework that systematically evaluates data sources across four dimensions: (i) sampled environmental spaces, (ii) model performance, (iii) predicted suitability patterns and (iv) model applicability and environmental extrapolation. By this, we evaluate model discrimination and applicability rather than predictive success for future invasion spread. We used the federal state of Hesse (Germany) as a case study, as it represents a well-monitored central European region where both CS and StAg data are sufficiently available to enable a comparative assessment. Specifically, we answer three research questions:

1. How does StAg and CS data affect the inferred environmental space of invasive plants in the invaded range of Hesse?
2. How do models based on StAg, CS and combined data differ in predictive performance and predicted habitat suitability for invasive plants in Hesse?
3. How does the data source affect the area of applicability (AOA) of the models, and along which environmental gradients are the training data underrepresented?

We expect inferred environmental spaces to differ between datasets, reflecting the observation processes the data is based on. Depending on the ecology of the species in place, we expect the integration of StAg and CS to a combined dataset to improve the performance, leading to broader environmental suitability predictions. Analogously, we expect a more complete area of applicability in the combined model compared to the single models.

## 2. Methods

### 2.1 Model species

To analyse differences between StAg and CS occurrence data, we initially considered all terrestrial invasive plants listed as of Union Concern recorded in Hesse (Study area description in Supplementary material 1), as these taxa generally have more extensive occurrence records and high management relevance due to EU legislation. For each candidate species, we queried occurrence counts in GBIF and requested StAg occurrence data from the Hessian State Agency for Nature Conservation, Environment, and Geology (HLNUG 2025d). Only those species with > 50 occurrences in both datasets were retained for our analysis, allowing for robust modelling (Hernandez et al., 2006; Soultan and Safi, 2017). This resulted in three model species: *Impatiens glandulifera* Royle, *Heracleum mantegazzianum* Sommier & Levier, and *Ailanthus altissima* (Mill.) Swingle (Supplementary material 1). *A. altissima* is a thermophilic tree species primarily occurring in warmer, urbanized areas, and its spreading process in Europe is expected to accelerate under climate change (HLNUG 2024; Pérez et al., 2022; Sladonja et al., 2015). The perennial monocarpic herb *H. mantegazzianum* is associated with linear corridors like roads, railways, or riverbanks, which also serve as dispersal pathways into surrounding habitats (Thiele et al., 2008). *I. glandulifera*, the most widespread of the three species, is an annual herb primarily dispersed via water and anthropogenic vectors. It is established both in linear structures and moist fallows or forests (Čuda et al., 2020; HLNUG 2024; Kiełtyk and Delimat, 2019).

### 2.2 Occurrence datasets

For each species, we used one global occurrence dataset and three regional datasets, each representing a different data source: citizen science (CS), state agency records (StAg), and a combined dataset. Occurrences records for the global model and the regional CS models were obtained from GBIF (2025a, 2025b). CS datasets originated from various publishers, including citizen science platforms like iNaturalist (inaturalist.org). StAg occurrences were sourced from the Hessian Agency for Nature Conservation, Environment and Geology (HLNUG 2025d; maps in Supplementary Material 1). The StAg dataset consists largely of data gathered through mapping performed as part of the “Hessische Biotopkartierung” (HB) from 1992 to 2006 and since 2014 the “Hessische Lebensraum-und Biotopkartierung” (HLBK), but also contains few occurrences reported by citizen scientists using the portal “Arten melden” (ca. 3 % of all occurrences in the StAg dataset; www.hlnug.de/themen/naturschutz/tiere-und-pflanzen/arten-melden). The goal of habitat mappings, which are carried out in sections of the study area at a time and executed by experts, is to map habitats of high conservation importance and also include information on invasive species in mapped habitats or surrounding areas. Each dataset thus carries its own biases, reflecting the distinct sampling processes and objectives behind their collection which is why invasion stage and survey period cannot be completely disentangled. The StAg data can be requested directly from HLNUG via the website (HLNUG 2025d). Finally, the CS and StAg occurrences were merged into a combined dataset.

For each species and data source, occurrences were filtered for year of record (≥ 1990) to match temporal scales of both datasets and coordinate uncertainty to match environmental data used for modelling (<1000 m). We further cleaned the datasets using the CoordinateCleaner package in R (Zizka et al., 2019) to identify and remove common spatial errors typically associated with CS data. To reduce spatial sampling bias and spatial autocorrelation, we additionally applied spatial thinning, retaining a single occurrence per 1 km^2^ raster cell (Beck et al., 2014; Boria et al., 2014; Supplementary material 2). The final occurrence numbers in the regional datasets were 2211 (CS) and 3999 (StAg) for *I. glandulifera*, 318 (CS) vs. 1427 (StAg) for *H. mantegazzianum* and 484 (CS) vs. 94 (StAg) for *A. altissima* (Supplementary material 2).

### 2.3 Environmental variables

As global climatic predictors, we chose the 19 bioclimatic variables from the CHELSA dataset (Brun et al., 2022) and kept the resolution of 1 km² for modelling as a tradeoff between spatial accuracy and computational requirements. We analysed the correlation between the variables and filtered out those with a Spearman correlation coefficient of |r| ≥ 0.7 (Dormann et al., 2013), retaining the more ecologically relevant variables based on species-specific considerations in line with literature (Supplementary material 3). This resulted in 6 climatic variables as global predictors: Annual Mean Temperature (Bio1), Temperature Seasonality (Bio4), Precipitation Seasonality (Bio15), Precipitation of Wettest Ǫuarter (Bio16), Precipitation of Driest Ǫuarter (Bio17), and Precipitation of Warmest Ǫuarter (Bio18).

For the regional habitat suitability models, we included climatic (Wan et al., 2015), topographic (BKG, 2021), infrastructure (BKG, 2024; HVBG, 2025, Weiss et al., 2018), hydrologic (BfG, 2024; HLNUG, 2025a; HLNUG, 2025c; HVBG, 2025), land use (EEA, 2019), and soil variables (HLNUG, 2025b; Supplementary Material 3). The regional variables were rasterized and standardized to a resolution of 1 km², spatially aligned with the global predictor grids to ensure consistency across model scales. Consistent with the global model, correlated predictors were filtered by retaining the ecologically more relevant variable, which resulted in 14 regional predictors (Supplementary Material 3).

### 2.4 Sampled environmental space

To analyse differences in sampled environmental space between CS and StAg data, we applied a PCA for all of Hesse and projected occurrences of both datasets into the PCA space. The environmental space was built using all numerical regional variables representing the topography, infrastructure, and hydrology. To aid in visualisation and exclude outliers, we projected a convex hull on the 95^th^ quantile for each data source. We then followed the workflow by Broennimann et al. (2012) to calculate density grids representing the environmental occupancy of each data source, and to compare the overlap of sampled environmental spaces. This method projects PCA scores of both the background environment and the species occurrences onto a regular grid in the two-dimensional PCA space. Occurrence points are then smoothed using a kernel density function, generating a continuous density surface. The resulting density grid reflects the local concentration of occurrences within the environmental space of each grid cell. From the density grids, we computed the quantitative overlap metric Schoener’s D (Schoener, 1970), ranging from zero to one, and evaluated the statistical significance of differences in environmental representation using the niche equivalency test of Warren et al. (2008). The test compares observed overlaps in environmental space to a null distribution generated by randomizing species occurrences (CS and StAg) within the PCA-projected environmental space of Hesse, while keeping the environmental background fixed. The null hypothesis states that differences in environmental space inferred from both datasets are not greater than expected by chance. To test this, occurrences records from both datasets are pooled and randomly reassigned between the two groups while maintaining original sample sizes and niche overlap is recalculated for each permutation. The observed Schoener’s D value is then compared against this null distribution.

### 2.5 Habitat suitability models

To assess the effect of data source on model outcome, we fitted separate models for each data source (CS, StAg, combined) following the hierarchical two-step modelling approach proposed by Gallien et al. (2012). We first modelled the global climatic suitability for our model species with the selected climatic variables. To minimize sampling bias and constrain model extrapolation to unobserved areas, pseudo-absences for the global models were randomly selected within a 1-20 km buffer surrounding known presences to restrict the absences to reachable locations (Gallien et al., 2012, Pereira et al. 2023). The number of pseudo-absence points was set equal to the number of presence records in each model, following recommendations for boosted regression tree algorithms (Barbet-Massin et al., 2012). The models were calculated using a generalized boosted regression tree algorithm with five iterations per species. Boosted regression trees were tuned using a grid search over a defined set of hyperparameters to identify the best-performing configuration for each iteration. In each iteration, 80% of the data was used for training and 20% for validation, with model training incorporating a 5-fold cross-validation.

We then continued modelling the regional habitat suitability for Hesse. For pseudo-absence selection, we used a combined approach of geographical and environmental constraints, since both have been shown to improve model performance (Barbet-Massin et al., 2012; Iturbide et al., 2015). Pseudo-absences were generated separately for each species × data source combination, following an identical workflow for all data sources. Occurrences of the respective dataset were projected into the environmental PCA space of Hesse, and only cells within the 0.55 quantile of occurrence distances to the occurrence centroid were retained as similar (environmental constraint). For the widespread *I. glandulifera*, the 0.95 quantile was used to ensure a sufficiently large selection space, as its occurrences cover most of the environmental space of Hesse (Section 3.1). Furthermore, only cells between 1 km and 20 km from occurrences of the respective dataset (geographical constraint) were retained as pseudo-absence selection space. This approach minimizes extrapolation into unobserved environmental space, thereby reducing the risk of unfounded predictions. The regional pseudo-absences were then inversely weighted based on predicted suitability from the global model, giving higher weight to low suitability regions, which leads to a more accurate selection of absences (Gallien et al., 2012). The number of pseudo-absences equalled the number of presences of the respective dataset and was redrawn in each iteration; as the global model was shared by all regional models, weights did not differ between data sources. At the regional scale, ten iterations were conducted to account for the smaller sample size and the resulting higher potential for model instability and imprecision compared to the global scale (Hernandez et al., 2006; Soultan and Safi, 2017). To avoid overfitting through autocorrelation of the data, we applied spatial block cross-validation (Meyer et al., 2019; Roberts et al., 2017). To this end, we partitioned the occurrences into six spatial blocks, one block being withheld as an independent validation set and the remaining five blocks used for cross-validation. Occurrences located within the extent of Hesse were excluded from the global model. Model performance for both the global and regional models was evaluated using the Area Under the Receiver Operator Curve (AUC; Phillips et al., 2006) and the true skill statistics (TSS; Allouche et al., 2006). Differences in AUC and TSS between data sources across iterations were tested with beta regressions (glmmTMB package Brooks et al. 2026; logit link), including species as a fixed effect, using likelihood-ratio tests for the effect of data source and its interaction with species, followed by Tukey-adjusted pairwise contrasts within species. Model assumptions were checked with simulated residuals (DHARMa package, Hartig and Lohse, 2022).

### 2.6 Analysis of suitability and area of applicability

#### 2.6.1 Predictor importance and response

To identify which environmental variables drive habitat suitability, we extracted variable importance and variable response for each data source and for each species.

#### 2.6.2 Area of applicability

We performed an AOA analysis for each species using the CAST package (Meyer and Pebesma, 2021). The AOA is calculated by applying a threshold to the dissimilarity index (DI), which quantifies the environmental distance between the prediction and the closest training data, with variables weighted by model importance. Dissimilarity should be interpreted as extrapolation-related uncertainty rather than total model uncertainty, as it does not differentiate between dataset specific biases and ecologically meaningful gradients. The AOA analysis was computed separately for each of the ten model iterations. To obtain an aggregated AOA raster, we applied a majority voting approach, considering a raster cell applicable if it fell within the AOA in at least eight out of the ten iterations. We explored which predictor ranges drive extrapolation beyond the AOA and how these differed between data sources. To achieve this, we first compared the number of cells classified as applicable by the averaged AOA.

We then fitted, for each species, data source and model iteration, an ordinary least-squares multiple linear regression of the DI on all predictors of the respective model, using only cells outside the AOA of that iteration. All predictors were entered jointly, so that coefficients represent conditional effects. Predictors were used in the form in which they entered the boosted regression tree model: continuous predictors were z-standardised with the mean and standard deviation of the training data (presences and pseudo-absences) of the respective iteration, and categorical predictors (CORINE land cover, floodplains, heat index, water-holding capacity) were dummy-coded with the first class as reference (CORINE: urban fabric; floodplains: class 1; heat index: “persistently hot”; water-holding capacity: water bodies) and standardized in the same way. As the DI is dimensionless, coefficients express the change in DI per standard deviation of a predictor in the training data of the respective model; for categorical predictors, per standard deviation of the class indicator. Because standardisation and DI both refer to the training data of each model, coefficients describe how strongly dissimilarity increases along each predictor relative to the environmental range covered by the respective dataset. Coefficients were averaged per species and data source across the iterations and also across species (unweighted mean). We then show predictors whose mean importance weight in the DI calculation exceeded the median across all predictors and data sources (Figure 3). To summarize the overall strength of the relationship per data source, we calculated the mean absolute coefficient across all predictors (each class of categorical predictors counted separately) per species and data source, and averaged it across species.

#### 2.6.3 Suitability patterns

To compare the proportion of applicable cells predicted as suitable between data sources, we converted the continuous suitability maps into binary representations using the threshold that maximizes the sum of sensitivity and specificity (Jiménez-Valverde and Lobo, 2007; Liu et al., 2016) and aggregated them into a consensus binary suitability map using the same majority voting approach as for the AOA maps. We then calculated the percentage of cells within each model’s AOA that were classified as suitable. As the AOA differs between models, these percentages refer to different reference areas. Next, we examined how the spatial extent and patterns of suitable areas within the applicable area varied across data sources. To this end, we extracted continuous suitability maps inside the AOA, averaged across 10 iterations, and quantified the overlap of these maps between data sources in a PCA-derived environmental space using Schoener’s D.

## 3. Results

### 3.1 Sampled environmental space

Variations in data sources led to a differentiation in the environmental representation of different species (Figure 1). The first PCA component explained approximately 27 % of the variance for all species, and the second component explained between 12.7% to 18.2 % of variance. Across species, PC1 represents the gradient from more remote, higher-elevation sites to lower, more urbanized areas. PC2 varies more among species but generally captures the gradient from higher-elevation or more urbanized sites towards wetter habitats located further from infrastructure. For all species, environmental spaces inferred from CS data are bound towards lower elevations and more urbanized or infrastructure-dominant areas. In contrast, StAg inferred environmental spaces oriented towards higher-elevation and low-infrastructure environments. This pattern is most pronounced for *A. altissima*, where each dataset captures substantial portions of the environmental space not represented by the other (Figure 1a). Across species, StAg inferred environmental spaces also showed a tendency to be located closer to watercourses compared to the CS inferred environmental spaces. In the PCA ordinations, the degree of the overlap between CS and StAg decreases from *I. glandulifera* > *H. mantegazzianum* > *A. altissima*. This is also represented in Schoener’s D values, ranging from 0.743 to 0.292. Warren’s test indicated significant differences between CS and StAg inferred environmental spaces for *A. altissima* and *I. glandulifera*, whereas no significant difference was found for *H. mantegazzianum*.

**Figure 1:**
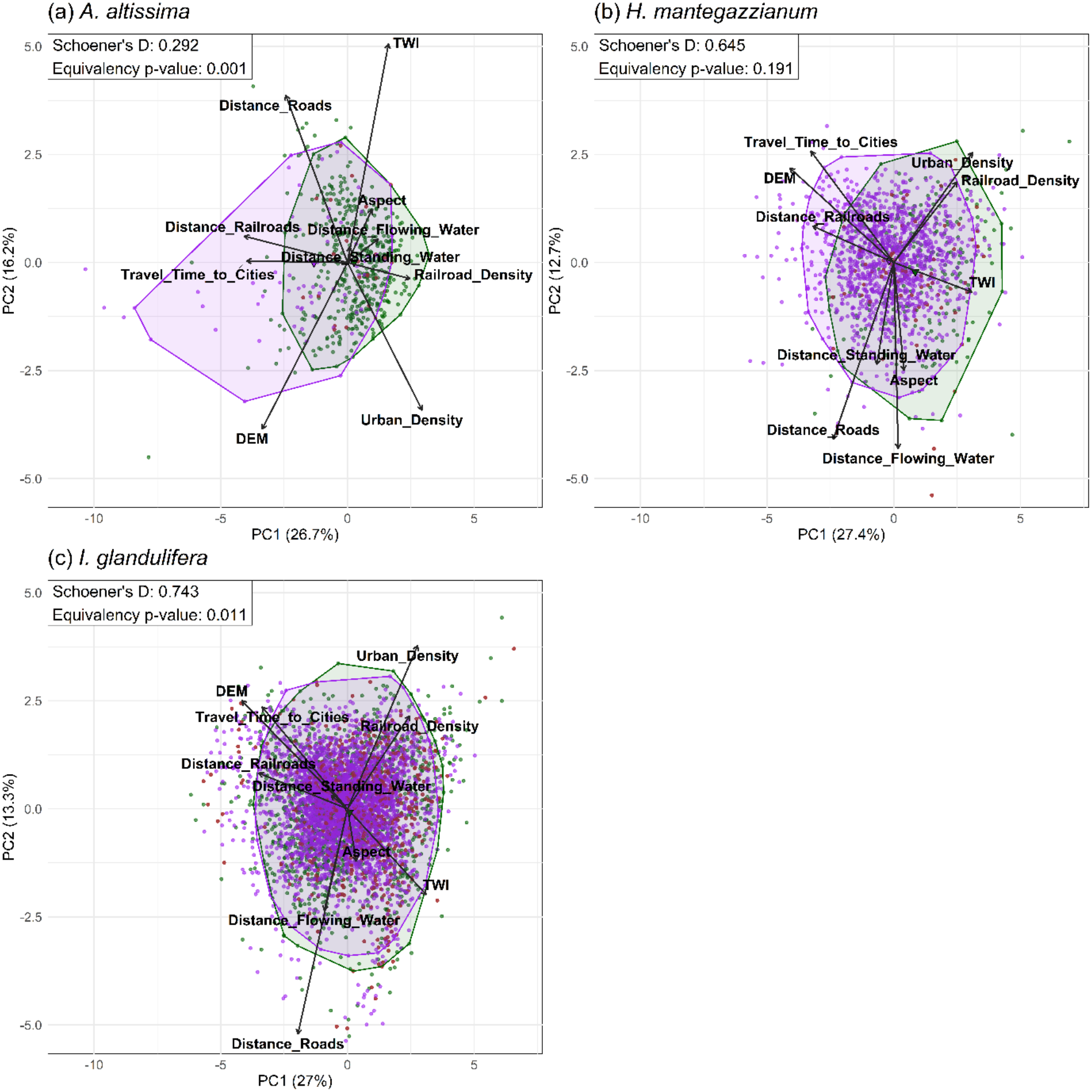
Environmental space sampled by citizen science (CS) and state agency (StAg) occurrence records of three invasive non-native plant species in the federal state of Hesse, central Germany: (a) *Ailanthus altissima*, (b) *Heracleum mantegazzianum*, (c) *Impatiens glandulifera*. The principal component analysis (PCA) was computed on all numerical regional predictors across Hesse (1 km × 1 km cells); axes show the first two principal components (PC1, PC2; dimensionless scores), with the percentage of variance explained in parentheses. Points are occurrences (green = CS, purple = StAg); polygons are convex hulls enclosing 95% of each source’s occurrences. Arrows show predictor loadings: DEM = elevation from the digital elevation model (m a.s.l.); Aspect (°); TWI = topographic wetness index (dimensionless); Travel_Time_to_Cities = travel time to the nearest city (min); Urban_Density = urban build-up density (0-1); Railroad_Density (m km⁻¹); Distance_Roads, Distance_Railroads, Distance_Flowing_Water, Distance_Standing_Water = distance to the nearest road, railroad, watercourse and standing water body (m). Boxes give Schoener’s D (dimensionless overlap of CS and StAg environmental spaces, from 0 = none to 1 = complete) and the p-value of Warren’s niche equivalency test (p < 0.05: spaces differ more than expected by chance).

### 3.2 Model performance

Across all species and data sources, models showed good discrimination, with AUC values ranging from 0.76 to 0.95 and TSS values from 0.42 to 0.81 (see Supplementary material 4). Averaged across species, the combined models yielded the highest mean AUC (mean = 0.85 ± sd = 0.07) and TSS (0.58 ± 0.15), followed by CS models (AUC = 0.83 ± 0.1; TSS = 0.57 ± 0.19) and StAg models (AUC = 0.82 ± 0.05; TSS = 0.54 ± 0.11). However, these differences were small, and after accounting for species identity there was no significant overall effect of data source on model performance across all iterations (beta regression, likelihood-ratio test; AUC: χ²(2) = 2.80, p = 0.25; TSS: χ²(2) = 2.65, p = 0.27). Instead, the effect of data source depended strongly on the species (source × species interaction; AUC: χ²(4) = 49.5, p < 0.001; TSS: χ²(4) = 36.9, p < 0.001; Supplementary Material 4). For *A. altissima* CS models performed best (AUC = 0.95 ± 0.02; TSS = 0.81 ± 0.05), and, together with combined models, significantly outperformed StAg models (pairwise contrasts, p < 0.01). For *H. mantegazzianum*, data sources did not differ significantly overall (AUC: χ²(2) = 5.58, p = 0.06; TSS: χ²(2) = 1.80, p = 0.41), with the highest AUC for combined models (0.80 ± 0.02) and the highest TSS for CS models (0.49 ± 0.09). For *I. glandulifera*, StAg models performed best (AUC = 0.83 ± 0.02; TSS = 0.52 ± 0.03), followed by combined and CS models, with all pairwise differences being significant (p < 0.05).

### 3.3 Predictor importance

Across the three data sources, topographic predictors consistently explained the most variance, followed by infrastructure and hydrology, together accounting for > 85 % of the mean model importance (Figure 2). Although their contributions remained consistent across all sources, some differences were apparent. CS models had a higher importance on infrastructure predictors, while StAg models were more explained by hydrology predictors, with combined models falling between the two. Response curves (see Supplementary material 5) also showed similar suitability responses to the predictor ranges of the key predictors. Suitability was generally similar in relation to elevation, but CS models showed higher suitability in lowlands compared to StAg models. Furthermore, CS also exhibited reduced suitability at greater distances from or lower density of infrastructure and a higher suitability further away from watercourses than COM and StAg models.

**Figure 2:**
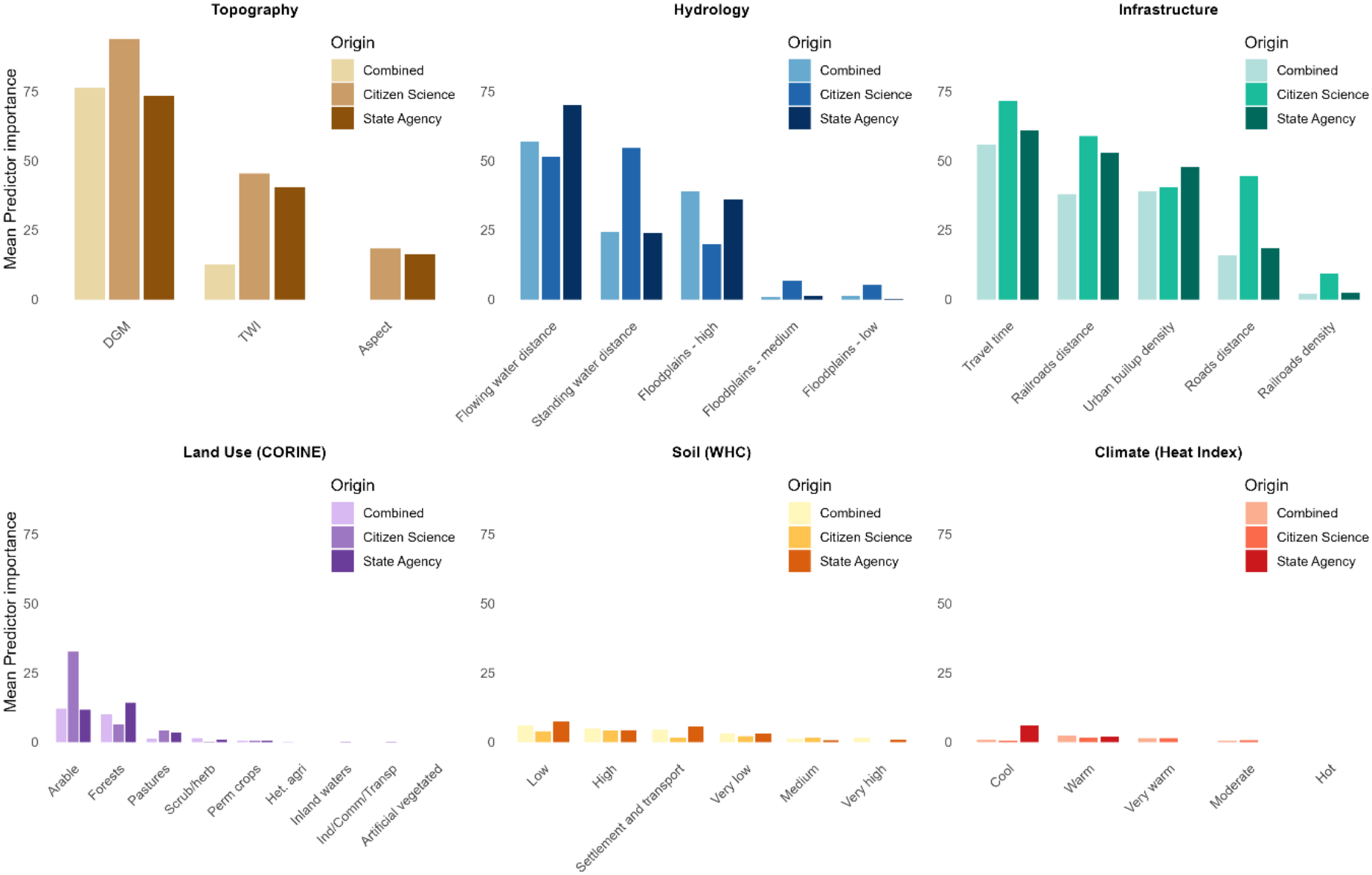
Mean relative importance of environmental predictors in the regional boosted regression tree models for Ailanthus altissima, Heracleum mantegazzianum and Impatiens glandulifera in the federal state of Hesse, central Germany, built from citizen science, state agency or combined (both sources pooled) occurrence data. Importance is dimensionless, scaled from 0 to 100 relative to the most influential predictor of each model, and averaged across 10 model iterations and the three species. Panels group predictors by category; categorical predictors are shown per class. DEM = elevation from the digital elevation model; TWI = topographic wetness index; CORINE = CORINE Land Cover classes (Het. agri = heterogeneous agricultural areas; Perm crops = permanent crops; Scrub/herb = scrub and/or herbaceous vegetation; Ind/Comm/Transp = industrial, commercial and transport units); WHC = soil water-holding capacity class; Heat Index = summer heat class (2001–2020). The Heat Index was used only for *A. altissima*, WHC only for *H. mantegazzianum* and *I. glandulifera*. Units of all predictors are given in Supplementary Material 3; response curves are in Supplementary Material 5.

### 3.4 Model applicability and environmental extrapolation

Spatial applicability was generally high across data sources but varied among species. All data sources showed a similar and high percentage of applicable cells for *I. glandulifera* and *H. mantegazzianum*, while for *A. altissima*, applicability was noticeably lower, with StAg models showing a higher applicability, followed by combined and CS models (Figure 4b).

Within non-applicable areas, environmental dissimilarity increased most strongly with elevation, urbanization predictors, and water-body-related predictors across data sources, as indicated by coefficients of the multiple regressions of DI on the predictors (Figure 3). The direction of these relationships was largely consistent across data sources, with the largest positive coefficients for the predictors along which the prediction area deviated most from the training data. Although dissimilarity increased along similar environmental gradients in all sources, the increase was consistently weaker in combined models (mean absolute coefficient = 0.017) than in single-source models (CS = 0.032; StAg = 0.031). Among the single-source models, dissimilarity in CS models increased more strongly at high elevations and with longer travel times to cities, whereas dissimilarity in StAg models increased more strongly with distance from watercourses. In addition, both StAg and combined models showed strong increases in dissimilarity in areas with a pronounced flooding regime (10-year flood discharge). At the species level, combined models also showed weaker increases in dissimilarity than single-source models for all species (Supplementary Material 6). Overall, coefficients were smallest for *I. glandulifera* (combined = 0.006; CS = 0.008; StAg = 0.008) and larger for *H. mantegazzianum* (combined = 0.013; CS = 0.041; StAg = 0.030) and *A. altissima* (combined = 0.033; CS = 0.047; StAg = 0.054). Species-specific patterns broadly followed the overall gradients, with two exceptions: for *I. glandulifera*, dissimilarity increased with proximity to cities, and for *H. mantegazzianum*, dissimilarity increased most strongly with distance from linear structures. *A. altissima* showed the largest coefficients across several predictors.

**Figure 3:**
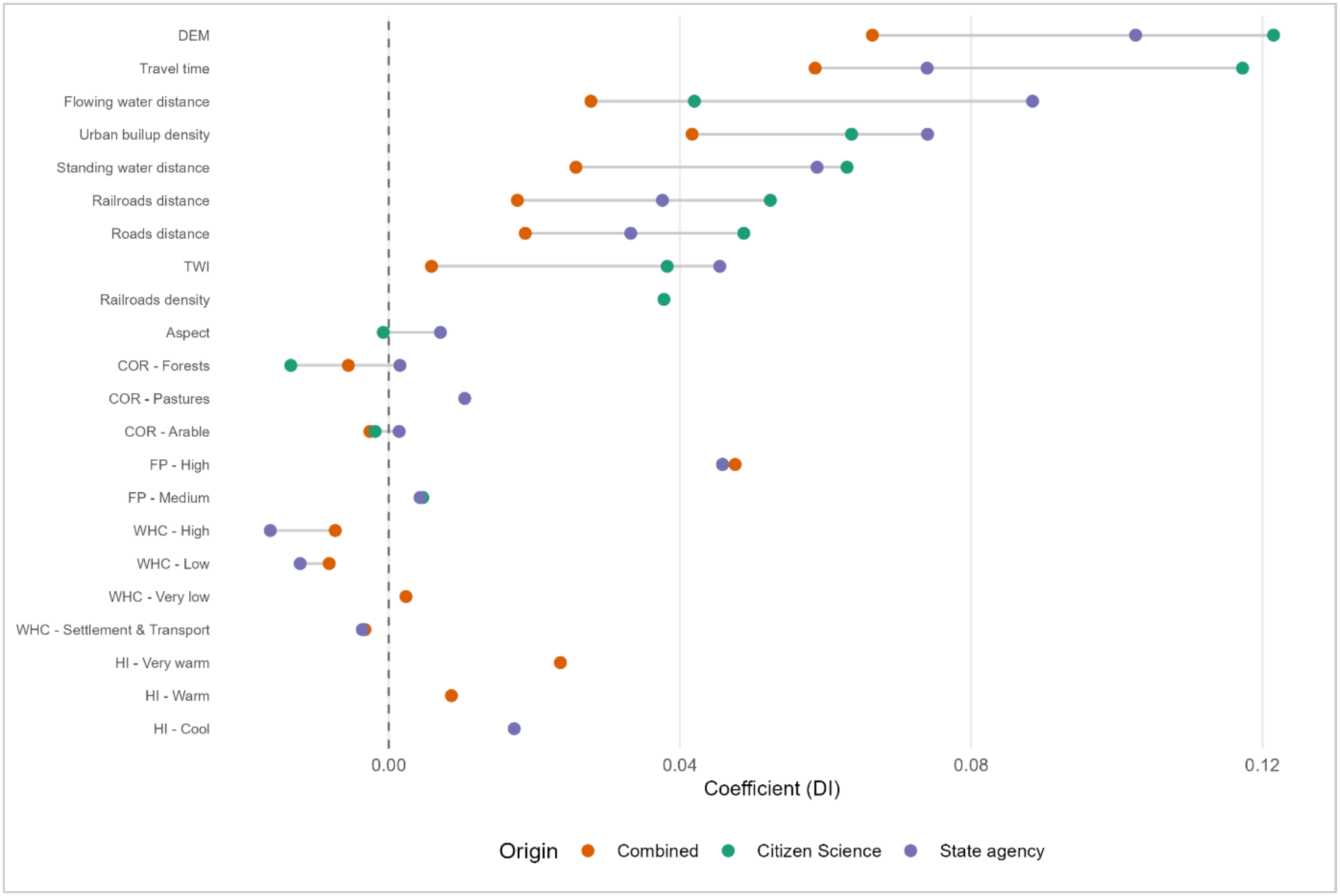
Environmental gradients driving extrapolation in the regional species distribution models for *Ailanthus altissima*, *Heracleum mantegazzianum* and *Impatiens glandulifera* in the federal state of Hesse, central Germany, built from citizen science, state agency or combined occurrence data. Points are standardized coefficients from multiple linear regressions of the dissimilarity index (DI) on all predictors, fitted to 1 km × 1 km cells outside the area of applicability (AOA). DI is the dimensionless, importance-weighted distance in predictor space between a cell and the nearest training data; cells whose DI exceeds a threshold derived from cross-validation lie outside the AOA, where the model extrapolates beyond the environmental conditions of the training data. Coefficients are dimensionless: the change in DI per standard deviation of a continuous predictor, or, for categorical predictors, per standard deviation of the class indicator (reference classes in Section 2.6.2). Positive values mean DI increases with higher predictor values; dashed line = zero; grey lines connect data sources. Only predictors with above-median importance are shown, so a missing point means below-median importance for that source. Values are averaged across 10 iterations and the three species (species-level results: Supplementary Material 6). DEM = elevation from the digital elevation model; TWI = topographic wetness index; COR = CORINE land-cover class; FP = floodplain class; WHC = soil water-holding capacity class; HI = heat index class (*A. altissima* only).

### 3.5 Suitability patterns

While the proportion of applicable cells predicted as suitable did not differ consistently between data sources (Figure 4a), suitability showed clear geographical differences between data sources (Figure 5). Within their respective AOA, combined models classified an intermediate or larger proportion of cells as suitable compared with CS and StAg models. Across data sources, this proportion was highest for *I. glandulifera*, followed by *H. mantegazzianum* and *A. altissima*. Because these proportions refer to the AOA of each model, they reflect both differences in predicted suitability and differences in the extent of the AOA (Figure 4b), and do not represent the total suitable area of Hesse.

**Figure 4:**
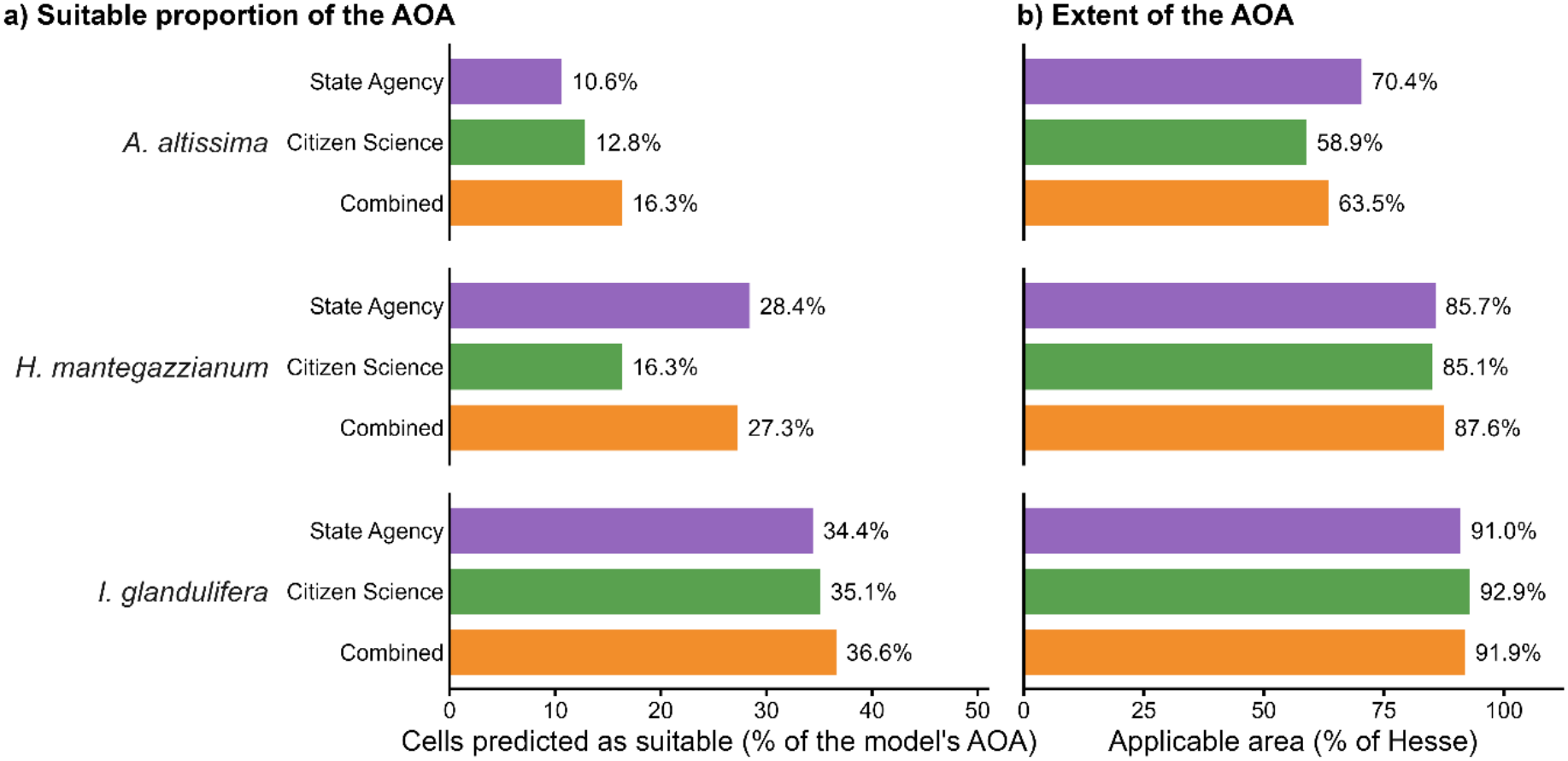
Area of applicability (AOA) and suitable proportion of the AOA of the regional species distribution models for *Ailanthus altissima, Heracleum mantegazzianum* and *Impatiens glandulifera* in the federal state of Hesse, central Germany, built from state agency, citizen science or combined occurrence data. (a) Percentage of 1 km × 1 km cells within each model’s AOA predicted as suitable; (b) percentage of Hesse within the AOA. The AOA comprises cells whose environmental conditions are similar enough to the training data (dissimilarity index, DI, below a threshold derived from cross-validation) for the cross-validated model performance to apply. Cells were counted as applicable or suitable if classified as such in ≥ 8 of 10 model iterations; continuous suitability was converted to suitable/unsuitable at the threshold maximizing the sum of sensitivity and specificity. Because the AOA differs between models, the percentages in (a) refer to different reference areas.

Concerning spatial suitability patterns, CS-based models consistently predicted higher suitability in southern and more urbanised regions of Hesse (Figure 5 d-f), whereas StAg models showed a spatial distribution extending into northern and rural areas while assigning lower suitability to suitability hotspots of CS models (Figure 5 g-i). combined models generally exhibited intermediate patterns, integrating patterns of both sources (Figure 5 a-c). This pattern was reflected in the Schoener’s D overlap analysis of suitability maps, with combined scoring higher similarity to both single-sourced models, while CS and StAg models showed lower overlap (Table 1).

**Figure 5:**
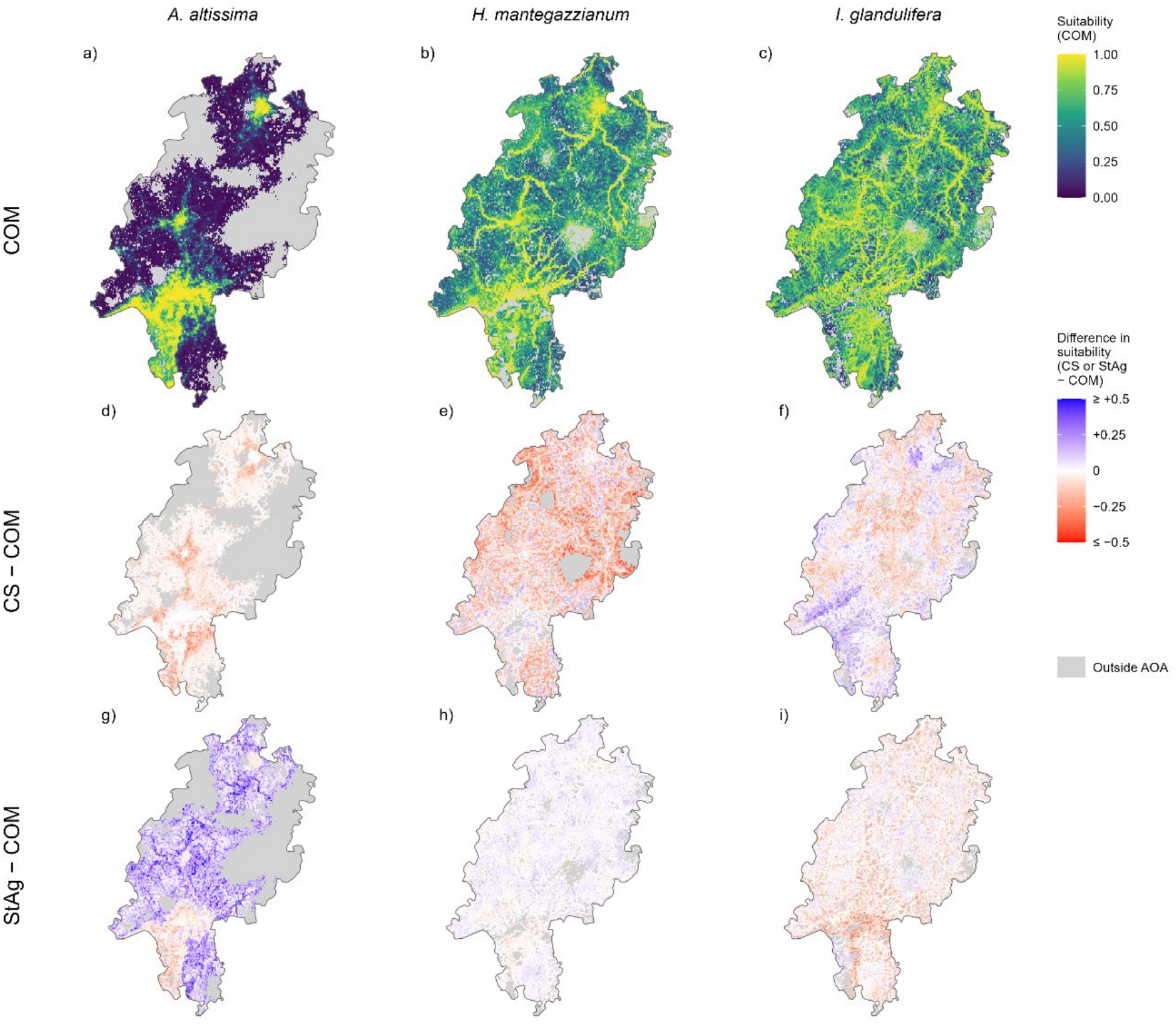
Predicted habitat suitability for Ailanthus altissima (a, d, g), Heracleum mantegazzianum (b, e, h) and Impatiens glandulifera (c, f, i) in the federal state of Hesse, central Germany, from regional boosted regression tree models averaged over 10 iterations. Top row (a– c): suitability from models based on combined citizen science and state agency data (COM). Suitability is dimensionless and ranges from 0 (unsuitable, dark purple) to 1 (highly suitable, yellow). Middle (d–f) and bottom (g–i) rows: cell-wise difference in suitability between the models based on citizen science data and combined (CS − COM) or between the models based on state agency data and combined (StAg − COM). Blue indicates higher suitability, red lower suitability and white no difference compared with the combined model. Observed differences range from −0.60 to +0.63; the color scale is truncated at ±0.5, and the few cells beyond this (0.08 %) are shown in the end colors. Predictions are shown only within the area of applicability (AOA), i.e. cells applicable in at least 8 of 10 iterations; cells outside the AOA have environmental conditions that differ too much from the training data. Differences (d–i) were computed only within the intersection of the AOAs of the two compared models, i.e. where both models are applicable. Grey cells lie outside the AOA of the combined model (a–c) or of at least one of the two compared models (d–i). Resolution 1 km × 1 km; WGS 84 (EPSG:4326).

**Table 1:** Pairwise Schoener’s D overlap of suitability-weighted environmental space between predicted suitability maps derived from different occurrence data sources (COM = Combined; CS = Citizen Science, StAg = State Agency).

| **Species** | **COM vs CS** | **COM vs StAg** | **CS vs StAg** |
|---|---|---|---|
| *I. glandulifera* | 0.954 | 0.962 | 0.928 |
| *A. altissima* | 0.906 | 0.757 | 0.680 |
| *H. mantegazzianum* | 0.886 | 0.959 | 0.872 |

Species-specific overlap was lowest for *A. altissima*, for which the suitability of CS models concentrated primarily in urbanized areas, whereas StAg models predicted suitability in a broader range of environments without pronounced clustering of high suitability areas (Figure 5). *H. mantegazzianum* showed more consistent suitability patterns across data sources, with combined predictions aligning more closely with StAg-based models than with CS-based models and being largely confined to river corridors. For *I. glandulifera*, the three models produced largely consistent predictions, with high suitability along rivers, and broader extensions into surrounding areas compared to *H. mantegazzianum*.

## 4. Discussion

By integrating state agency monitoring data with citizen science records, we show that the two data sources sample distinct, complementary parts of the environmental space of invasive plants, a difference that standard performance metrics may fail to reveal. Our analysis shows that integrating data sources alters suitability predictions and expands the environmental coverage of the training data, while effects on internal discrimination were small and species-dependent. This is due to the different sampling strategies datasets are based on: StAg data better captured remote, high-elevation areas often missed by CS and combined models provided a higher overlap and weaker increases in dissimilarity along environmental gradients. AOA analyses showed that combined models did not consistently improve spatial applicability but reduced dissimilarity along environmental gradients. This may be relevant for invasion risk assessments, where single-source models may under- or overestimate suitability in habitats critical for conservation, such as riparian zones.

### 4.1 Environmental representation is data source dependent

Our results highlight the importance of combining data sources with different sampling designs to represent the environmental space invaded by each species. StAg and CS occurrence data cover different parts of the environmental space occupied by invasive plants because they are collected under fundamentally different sampling frameworks: CS occurrences, which are often collected spontaneously through species identification applications (Klinger et al., 2023), were more strongly associated with lower-elevation, human-influenced environments. This aligns well with studies showing that opportunistically collected occurrences are spatially biased towards accessible locations, especially close to population centres and infrastructure (Geldmann et al., 2016; Mair and Ruete, 2016). StAg occurrences, which were collected based on efforts to map protected habitats, covered environmental spaces in higher-elevation, remote areas which are closer to watercourses compared to CS (Cazzolla Gatti et al., 2023; Joppa and Pfaff, 2009). To this end, StAg data is a suitable complement to CS data, as both datasets are biased because of the underlying sampling process. Thus, environmental spaces covered by each data source may represent different parts of the full realized environmental space, with neither dataset capturing its full extent. Differences in sampled environmental space between CS and StAg also differed between species, as they were largest for *A. altissima*, the species with the most restricted distribution in Hesse, and smallest for the widespread *I. glandulifera*. One possible explanation may be that the realized environmental space of less widespread species is small relative to the regional environmental context, so that different sampling processes capture different parts of it. In contrast, generalist species like *I. glandulifera* occupy a large part of the available environmental space in Hesse, leading to more similar estimates across different datasets. We hypothesize that the relevance of such differences also depends on ecological context, such as habitat preference (e.g., specialists vs generalists; Liu et al., 2020), invasion stage (e.g., early spread vs established; Berio Fortini et al., 2024) and pathways of spread, and that differences between datasets may be more pronounced for more recently introduced invasive plants. As our study includes only three species, these relationships remain tentative and testing them would require comparative analyses across a larger number of taxa. At the same time, these differences cannot be attributed to species ecology or invasion stage alone, as they also reflect the sampling objectives and geographical coverage of each observation process.

Overall, both CS and StAg data capture different biased subsets of the real species distribution, which is why combining them may increase environmental space representation due to the complementary sampling strategies. Combining two biased sources does not, however, remove their biases: in the combined dataset, areas missed by both sampling processes, such as remote, high-elevation sites, may remain underrepresented. Based on the available data, we cannot fully differentiate between sampling bias, true absences unsuitable environments and uncolonized suitable areas. This issue can only be resolved by performing structured monitoring for invasive plants that reports both presences and absences.

### 4.2 Impact on SDM prediction

SDMs calibrated with combined occurrences may have resulted in more extensive suitability patterns, which is consistent with previous studies integrating multiple occurrence sources into SDMs on plants (Crall et al., 2015; Dimson et al., 2023) and birds (Robinson et al., 2020). When comparing the CS and StAg models, differences in occurrence data were reflected both in the spatial suitability patterns and in the predictor importance. This is in line with other studies on invasive species modelling, where adding citizen data generally expanded known distributions, environmental gradients sampled, and made habitat suitability maps more realistic, even when AUC gains were small (César de Sá et al., 2019; Crall et al., 2015). In our study, Schoener’s D values indicated that SDMs reduce divergence between CS and StAg occurrences while not fully eliminating data-source-specific patterns. Like the data sets, CS models placed more importance on predictors associated with urbanisation and infrastructure whereas StAg models produced broader suitability estimates while showing higher importance in hydrological predictors and represented rural regions. Interestingly, all models achieved high discrimination performance across data sources. However, these metrics were not systematically related to ecologically meaningful suitability patterns, a limitation widely recognised in SDM literature (Bracho-Estévanez et al., 2024; Fourcade et al., 2018; Lobo et al., 2008). This was most evident for *A. altissima*, where models calibrated with CS data achieved the highest AUC scores, but failed to capture the environmental conditions covered by StAg data, resulting in high discriminatory ability for only a limited part of the species’ environmental distribution. Because each model was evaluated on test data from its own source, high AUC here reflects discrimination within the environmental space sampled by CS records, not across the conditions represented by StAg data. In general, AUC and TSS assess discrimination within our evaluation framework, but do not establish which model better represents the actual distribution of the species. In contrast, combined models rarely achieved the highest performance for individual species but yielded stable results when averaged across species. This pattern is consistent with a study showing that integrated data increased realized environmental coverage and led to higher predicted invasion risk, especially for terrestrial species, but for some aquatic species resulted in environmental space contraction and lower predicted risk (Di Febbraro et al., 2023). Overall, model results probably reflect the sampling processes of the underlying datasets as much as the ecological preferences of the study species, as both presences and pseudo-absences, drawn around each dataset’s records, inherit the spatial footprint of the respective observation process.

### 4.3 Environmental coverage, applicability and monitoring applications

Combining CS and StAg data into combined models did not consistently improve the area of applicability. This may either reflect constraints imposed by the species’ realized environmental space in the study region or be the result of dataset-specific biases, since the AOA threshold depends on the density distribution of environmental conditions covered in the training data (Meyer and Pebesma, 2021). For *A. altissima*, combined models showed higher spatial applicability than CS models, indicating that AOA can be enhanced when occurrences from different data sources cover different parts of the occupied environmental space. This may be because the *A. altissima* is spreading from cities to natural habitats (Lezcano Caceres, 2010), whereas *H. mantegazzianum* and *I. glandulifera* may have colonized a high proportion of suitable areas. DI analysis further showed that dissimilarity to the training data increased less strongly along environmental gradients in combined models than in single-source models, indicating that a larger part of the environmental conditions in Hesse lies closer to the represented environmental space. This is somewhat contrasting to the findings of Di Febbraro et al. (2023), where citizen science data negatively affected the environmental space quantification and the prediction of future biological invasion risk in some aquatic species. Despite this, remote and high-elevation areas remained outside or at the margin of the training domain in all models. This pattern likely reflects a combination of true absences and limited sampling effort in either dataset, as these areas are more distant from dispersal corridors for invasive plants such as road networks and water bodies (Hulme et al., 2008; Roy et al., 2024). Nonetheless, invasive plants often show declining occurrences with increasing elevation and are typically more abundant in lowlands influenced by human activity (Alexander et al., 2016; Vorstenbosch et al., 2020). This is partly due to climatic constraints, as observed for *A. altissima* (Motti et al., 2021; Sladonja et al., 2015), and partly due to reduced propagule pressure in less accessible areas such as mountains (Alexander et al., 2016). At the same time, invasive plants such as *I. glandulifera* have been reported to expand into higher elevations (Zając et al., 2011) where monitoring intensity tends to decline. Consequently, it is only partially possible to distinguish true absences from under-monitored areas. The AOA and dissimilarity analyses can therefore identify environmental conditions that are underrepresented in the training data, but they cannot determine whether this underrepresentation reflects insufficient survey effort, unsuitable habitat or environments that have not yet been colonized. In addition, spatial and temporal biases in the observations can limit their overall representativeness (Di Cecco et al., 2021).

Within these limits, the framework has several potential applications for monitoring and management. Integrating CS observations with institutional datasets highlights the complementarity between participatory approaches and more traditional monitoring schemes (Lozano et al., 2017). Environments that are underrepresented in the training data of all models, here remote, high-elevation sites and riparian corridors, could be used to prioritize targeted surveys. If such surveys record both presences and absences, they would also allow monitoring gaps to be distinguished from unsuitable or uncolonized environments. Our results further suggest that risk maps based on GBIF records alone may underestimate the potential distribution of spreading species such as *A. altissima* outside urban areas, while agency data alone may underrepresent urban source populations. Risk assessments and the prioritization of management for species of Union concern could therefore benefit from combining both data sources. Including remote-sensing data could further improve the representation of environmental space and model performance, as shown for *Prosopis juliffora* in Ethiopia (Ahmed et al., 2021). Future research could thus explore how to formally integrate citizen science into hybrid monitoring frameworks that combine institutional surveys and promote adaptive surveillance systems.

### 4.4 Conclusions

Our results indicate that integrating datasets with different sampling strategies in species distribution models can alter modelling results and could guide monitoring and management priorities. Integrating CS and StAg data expanded the environmental representation of occurrence records, while effects on internal discrimination were small and species-dependent. Our study can therefore serve as a transferable analytical workflow, not as proof of a universal performance advantage for combined datasets and the generality of these findings needs testing across more taxa and regions. Outcomes may depend on various factors such as invasion stage, or habitat specialization. Integrating datasets may be relevant in particular for species with narrow or patchy distributions, where both CS and StAg data may fail to capture the realized environmental space. To close potential monitoring gaps, state agencies could target environments underrepresented in both datasets, combined with the systematic documentation of absences. . Finally, authorities should make occurrence data available in accordance with the FAIR principles, as data availability improves early detection and coordinated responses to invasions.

## Data statement

Citizen science occurrence data are available from GBIF (Hesse: https://doi.org/10.15468/dl.865r6h; global: https://doi.org/10.15468/dl.ns4htj). The filtered occurrence tables used for modelling are included in the code repository. State agency occurrence data are available on request from the Hessian Biodiversity Database (HEBID) of the Hessian State Agency for Nature Conservation, Environment and Geology (HLNUG). The processed environmental predictor layers of the regional models are provided in the code repository together with their original sources, licenses and processing steps; the global bioclimatic predictors are available from CHELSA v2.1 (Brun et al., 2022). The R code used for all analyses is available at https://github.com/YvesKlinger/INNS_model_Hesse and archived at Zenodo https://doi.org/10.5281/zenodo.24103587.

## Acknowledgements

We are grateful to the citizen scientists that collected occurrence data that we used in this study as well as to the Hessian Biodiversity Database.

## Declaration of generative AI and AI-assisted technologies in the manuscript preparation process

During the preparation of this work, the authors used Claude Code (version 2.1.263; model: Claude Opus 5.5; Anthropic) to debug and cleanup analysis code. All code produced or altered with this tool was manually reviewed, tested and verified by the authors. After using this tool, the authors reviewed and edited the content as needed and take full responsibility for the content of the published article.

## Supplementary material

### Appendix 1.1: Study area description

The federal state of Hesse (ca. 21,116 km²; Statistisches Bundesamt, 2024) is located in central Germany and spans an elevation from ca. 75 m a.s.l. in the lowlands along the Rhine and Main rivers in the south to ca. 950 m a.s.l. in the Rhön in the east (NASA JPL, 2020). Forests cover ca. 42% of the state (HMLU, 2024), whereas the south is dominated by the densely populated and urbanized Frankfurt–Rhine/Main area. Forest cover is lowest in intensively farmed regions such as the Wetterau north of Frankfurt (ca. 15%; HMLU, 2024; Jauker et al., 2009), while peripheral low mountain regions such as the Lahn–Dill district in western Hesse contain large shares of extensively managed grassland (Reger et al., 2009). Hesse has a moderately continental climate. In most parts of the state, mean annual precipitation is between 600 and 800 mm, decreasing to ca. 500 mm in the Upper Rhine Valley and exceeding 1,400 mm in the highest parts of the Vogelsberg and the Rhön (HLNUG, 2018). Mean annual temperature is around 8.8 °C (1981–2010) and decreases from south to north (HLNUG, 2018; Huebener et al., 2022). Together, these patterns result in strong environmental gradients within Hesse, from the warm, urbanized lowlands in the south to the forest- and grassland-dominated low mountain ranges in the centre and north. These gradients are relevant for our model species, since *A. altissima* is mainly confined to warm, urbanized areas (Sladonja et al., 2015), whereas *H. mantegazzianum* and *I. glandulifera* spread along linear structures such as roads, railways and watercourses (Čuda et al., 2020; Thiele et al., 2008).

---

**Appendix 1.2:** Number of occurrence records per 1 km × 1 km cell of the three invasive non-native plant species *Ailanthus altissima*, *Heracleum mantegazzianum* and *Impatiens glandulifera* in the federal state of Hesse (central Germany) from citizen science (CS; top row) and state agency data (StAg; bottom row). Records were filtered for year of record (≥ 1990) and coordinate uncertainty and cleaned with the CoordinateCleaner package (Zizka et al., 2019) as described in Section 2.2 of the publication. Records are shown as numbers before spatial thinning and the number of occupied cells corresponds to the number of records retained after thinning (one record per cell). Colors indicate the number of records per cell in four classes (1, 2–3, 4–10, > 10); white areas contain no records. Each panel gives the number of records and of occupied cells. Maps are projected in ETRS89 / UTM zone 32N (EPSG: 25832).

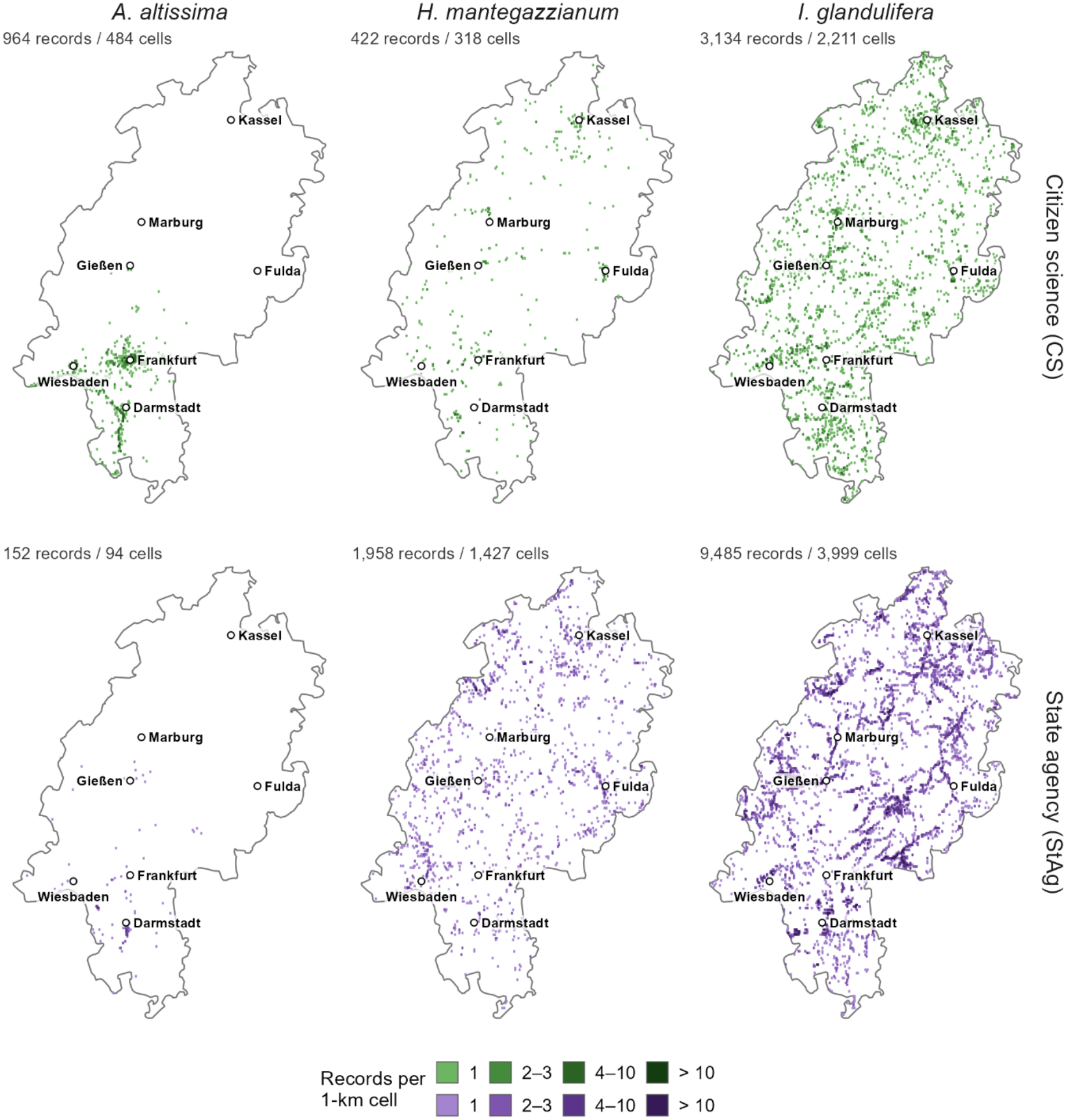

**Appendix 2:** Number of occurrence records per species and dataset before filtering, after filtering and coordinate cleaning, and after spatial thinning. The records after thinning were used for modelling.

| **Dataset** | **Step** | ***I. glandulifera*** | ***H. mantegazzianum*** | ***A. altissima*** |
|---|---|---|---|---|
| Global (GBIF)<sup>a</sup> | Before | 413060 | 148357 | 135284 |
|  | After filtering and cleaning<sup>c</sup> | 328470 | 107849 | 117974 |
|  | After thinning<sup>d</sup> | 119927 | 37871 | 54520 |
| Regional CS (GBIF)<sup>a</sup> | Before | 3677 | 510 | 1242 |
|  | After filtering and cleaning<sup>c</sup> | 3134 | 422 | 964 |
|  | After thinning<sup>d</sup> | 2211 | 318 | 484 |
| Regional StAg (HLNUG)<sup>b</sup> | Before | 9621 | 2120 | 162 |
|  | After filtering and cleaning<sup>c</sup> | 9485 | 1958 | 152 |
|  | After thinning<sup>d</sup> | 3999 | 1427 | 94 |
| Regional COM<sup>e</sup> | Before | 13298 | 2630 | 1404 |
|  | After filtering and cleaning<sup>c</sup> | 12619 | 2380 | 1116 |
|  | After thinning<sup>d</sup> | 5523 | 1662 | 553 |

<sup>a</sup> GBIF records were downloaded with the following filters already applied to the query: year of record ≥ 1990; coordinate uncertainty < 1000 m or not reported; occurrence status "present"; basis of record human observation, machine observation, observation or occurrence evidence; coordinates available and no geospatial issues flagged by GBIF. The regional download was further restricted to Hesse (GADM DEU.7\_1). "Before" therefore refers to the downloaded records, not to all GBIF records of the species (GBIF 2025a, 2025b).
<sup>b</sup> StAg records were filtered for year of record (≥ 1990) and positional accuracy. Records whose location was given only at grid-quadrant level or was flagged as imprecise were removed. This step left 9496, 1977 and 156 records for *I. glandulifera, H. mantegazzianum* and *A. altissima*, respectively, before coordinate cleaning.
<sup>c</sup> Coordinate cleaning used the CoordinateCleaner package (Zizka et al., 2019). It removed records with invalid coordinates, zero coordinates, identical latitude and longitude, coordinates at country or province centroids, at sea, at the GBIF headquarters or at biodiversity institutions, geographic outliers, and records duplicating the coordinates of another record of the same species.
<sup>d</sup> Spatial thinning kept one record per grid cell. The regional datasets used the 1 km × 1 km predictor grid of Hesse, and the global dataset used the 30 arc-second CHELSA grid (Brun et al., 2022). Before thinning the global dataset, records within the rectangular extent of Hesse were removed so that the global and regional calibration data are independent. The global "after thinning" count therefore also reflects this exclusion.
<sup>e</sup> The COM dataset was created by merging the CS and StAg records before filtering, and was then filtered and cleaned as described in <sup>b</sup> and <sup>c</sup>. Thinning was applied to the merged dataset. Cells occupied by records from both sources were therefore counted only once, which is why COM counts after thinning are lower than the sum of CS and StAg.

**Appendix 2.1:** Reasoning for excluding and retaining variables

| Species affected | Variables excluded | Variables retained | Reasoning (see also Appendix 2) |
|---|---|---|---|
| All | All other bioclimatic variables (Brun et al. 2022) | Bio1, Bio4, Bio15, Bio16, Bio17, Bio18 (Brun et al. 2022) | Key variables describing climatic means and extremes constraining distribution. |
| *A. altissima* | WHC (HLNUG, 2025) | Heat index (Wan et al. 2015) | Temperature variables capturing, e.g., urban heat effects, were included to account for the preference for warm climates. |
| *H. mantegazzianum & I. glandulifera* | Heat index (Wan et al. 2015) | WHC (HLNUG, 2025) | Water-related variables were prioritized over temperature variables, as water availability and drought stress are expected to be the main limiting factors. |

**Appendix 3.2:** Environmental variables retained for SDMs after correlation analysis (|r| ≥ 0.7) according to their use in the Global or the Local models and categorized in Climate, Topography, Infrastructure, Hydrology, Land use, and Soil variables. A brief reasoning for excluding and retaining variables in the models can be found in Supplementary Material 3.1.

| Model Extent | Category | Variables | Units | Dataset Reference | Reasoning / Reference |
|---|---|---|---|---|---|
| Global | Climate | Annual Mean Temperature (**Bio1**),<br>Temperature Seasonality (**Bio4**),<br>Precipitation Seasonality (**Bio15**),<br>Precipitation of Wettest Quarter (**Bio16**),<br>Precipitation of Driest Quarter (**Bio17**),<br>Precipitation of Warmest Quarter (**Bio18**) | °C<br>°C/100<br>kg m<sup>-2</sup><br>kg m<sup>-2</sup> month<sup>-1</sup><br>kg m<sup>-2</sup> month<sup>-1</sup><br>kg m<sup>-2</sup> month<sup>-1</sup> | CHELSA v2.1 (Brun et al., 2022) | Captures broad-scale climatic conditions to capture species climatic niche (e.g., used in Gallien et al. 2012) |
| Local | Climate | **Heat Index**\* | categorical | Average heat index (summer months 2001-2020; Wan et al., 2015) | Represents local heat conditions, relevant for *A. altissima* since it is confined to heat islands (e.g., urban areas) outside its climatic niche (Sladonja et al. 2015) |
| Local | Topography | Digital Elevation Model (**DEM**)<br>**Aspect**<br>Topographic Wetness Index (**TWI**) | m a.s.l.<br>°<br>unitless | BKG (2021)<br>Calculated from DEM<br>Calculated from DEM | Topographic variables capture microclimates concerning radiation, precipitation, moisture, and air temperature |
| Local | Infrastructure | **Travel time to nearest city**<br>**Urban buildup density**<br>**Railroad density**<br>**Railroad distance**<br>**Road distance** | minutes<br>share of cell (0–1)<br>m * km<sup>-1</sup><br>m<br>m | Weiss et al. (2018)<br>BKG (2024), HVBG (2025)<br>BKG (2024), HVGB (2025)<br>BKG (2024), HVGB (2025)<br>BKG (2024), HVGB (2025) | Proxy for human influence and propagule pressure. |
| Local | Hydrology | **Standing water distance**<br>**Flowing water distance**<br>**Floodplains** | m<br>m<br>categorical | BfG (2024), HVBG (2025)<br>BfG (2024), HLNUG (2025a)<br>HLNUG (2025c) | Affects moisture availability and propagule pressure. |
| Local | Land use | **CORINE land cover** | categorical | EEA (2019) |  |
| Local | Soil | Water holding capacity (**WHC**)\*\* | categorical | HLNUG (2025b) | Relevant for *H. mantegazzianum* & *I. glandulifera* as they are sensitive to soil moisture availability. |

\* only for *A. altissima* models
\*\*only for *H. mantegazzianum* & *I. glandulifera* models

**Appendix 3.1:** AUC and TSS values across n=10 model iterations for each combination of data source and species in the regional habitat suitability models.

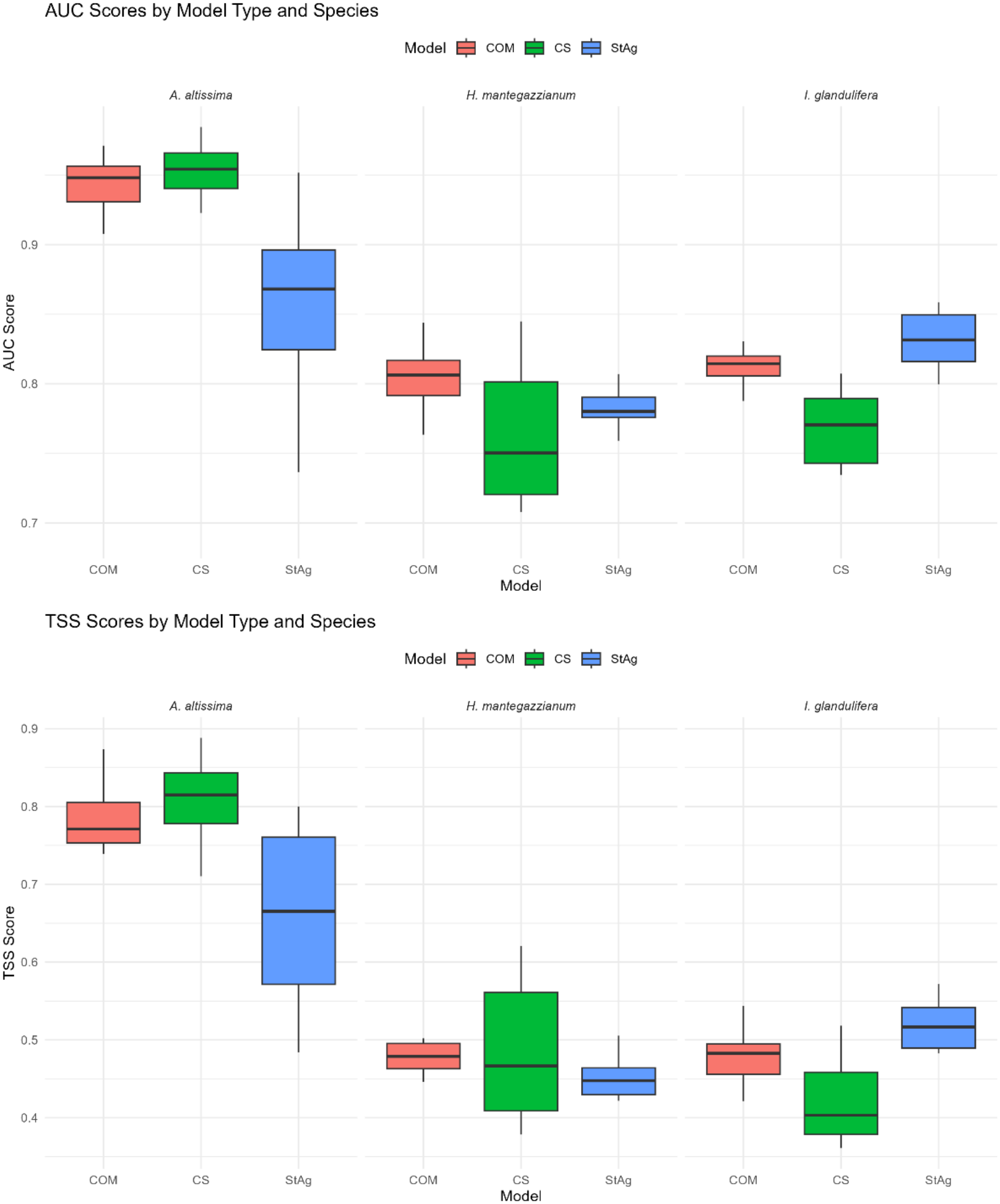

**Appendix 4.2:** Likelihood-ratio tests of differences in test performance (AUC, TSS) between data sources (COM, CS, StAg) of the local models across all model runs, based on beta regressions. Overall source effect: source + species vs. species only; source × species interaction: source × species vs. source + species; source effect within species: source vs. intercept-only model fitted separately per species (10 runs per data source and species). Significant p-values (< 0.05) in bold.

| Test | Metric | Species | n | χ<sup>2</sup> | df | p |
|---|---|---|---|---|---|---|
| Overall source effect | AUC | All species | 90 | 2.80 | 2 | 0.247 |
|  | TSS | All species | 90 | 2.65 | 2 | 0.266 |
| Source × species interaction | AUC | All species | 90 | 49.54 | 4 | **< 0.001** |
|  | TSS | All species | 90 | 36.87 | 4 | **< 0.001** |
| Source effect within species | AUC | *A. altissima* | 30 | 18.13 | 2 | **< 0.001** |
|  | AUC | *H. mantegazzianum* | 30 | 5.58 | 2 | 0.061 |
|  | AUC | *I. glandulifera* | 30 | 27.64 | 2 | **< 0.001** |
|  | TSS | *A. altissima* | 30 | 14.93 | 2 | **< 0.001** |
|  | TSS | *H. mantegazzianum* | 30 | 1.80 | 2 | 0.407 |
|  | TSS | *I. glandulifera* | 30 | 22.24 | 2 | **< 0.001** |

COM = combined data; CS = citizen science data (GBIF); StAg = state agency data (HLNUG).

**Appendix 4.3:** Pairwise differences in mean test performance between data sources within each species, estimated from the species-specific beta regressions (estimated marginal means on the response scale). Differences are given as first minus second data source; 95% confidence intervals and p-values are Tukey-adjusted. Significant p-values (< 0.05) in bold.

| Species | Metric | Contrast | Difference | 95% CI | z | p |
|---|---|---|---|---|---|---|
| *A. altissima* | AUC | COM – CS | -0.016 | [-0.051, 0.019] | -1.07 | 0.535 |
|  | AUC | COM – StAg | 0.076 | [0.029, 0.123] | 3.78 | < **0.001** |
|  | AUC | CS – StAg | 0.092 | [0.047, 0.137] | 4.78 | < **0.001** |
|  | TSS | COM – CS | -0.037 | [-0.109, 0.035] | -1.21 | 0.446 |
|  | TSS | COM – StAg | 0.106 | [0.026, 0.185] | 3.10 | **0.005** |
|  | TSS | CS – StAg | 0.143 | [0.065, 0.220] | 4.33 | < **0.001** |
| *H. mantegazzianum* | AUC | COM – CS | 0.036 | [0.002, 0.070] | 2.48 | **0.035** |
|  | AUC | COM – StAg | 0.019 | [-0.015, 0.052] | 1.30 | 0.398 |
|  | AUC | CS – StAg | -0.018 | [-0.052, 0.017] | -1.18 | 0.465 |
|  | TSS | COM – CS | -0.007 | [-0.067, 0.053] | -0.27 | 0.961 |
|  | TSS | COM – StAg | 0.026 | [-0.034, 0.086] | 1.02 | 0.564 |
|  | TSS | CS – StAg | 0.033 | [-0.027, 0.093] | 1.29 | 0.401 |
| *I. glandulifera* | AUC | COM – CS | 0.041 | [0.019, 0.064] | 4.31 | < **0.001** |
|  | AUC | COM – StAg | -0.021 | [-0.042, -0.000] | -2.36 | **0.048** |
|  | AUC | CS – StAg | -0.062 | [-0.084, -0.040] | -6.67 | < **0.001** |
|  | TSS | COM – CS | 0.059 | [0.018, 0.100] | 3.36 | **0.002** |
|  | TSS | COM – StAg | -0.042 | [-0.083, -0.000] | -2.37 | **0.047** |
|  | TSS | CS – StAg | -0.100 | [-0.141, -0.059] | -5.74 | < **0.001** |

COM = combined data; CS = citizen science data (GBIF); StAg = state agency data (HLNUG).

**Appendix 5.1:** Response plots for all used predictors per data source.

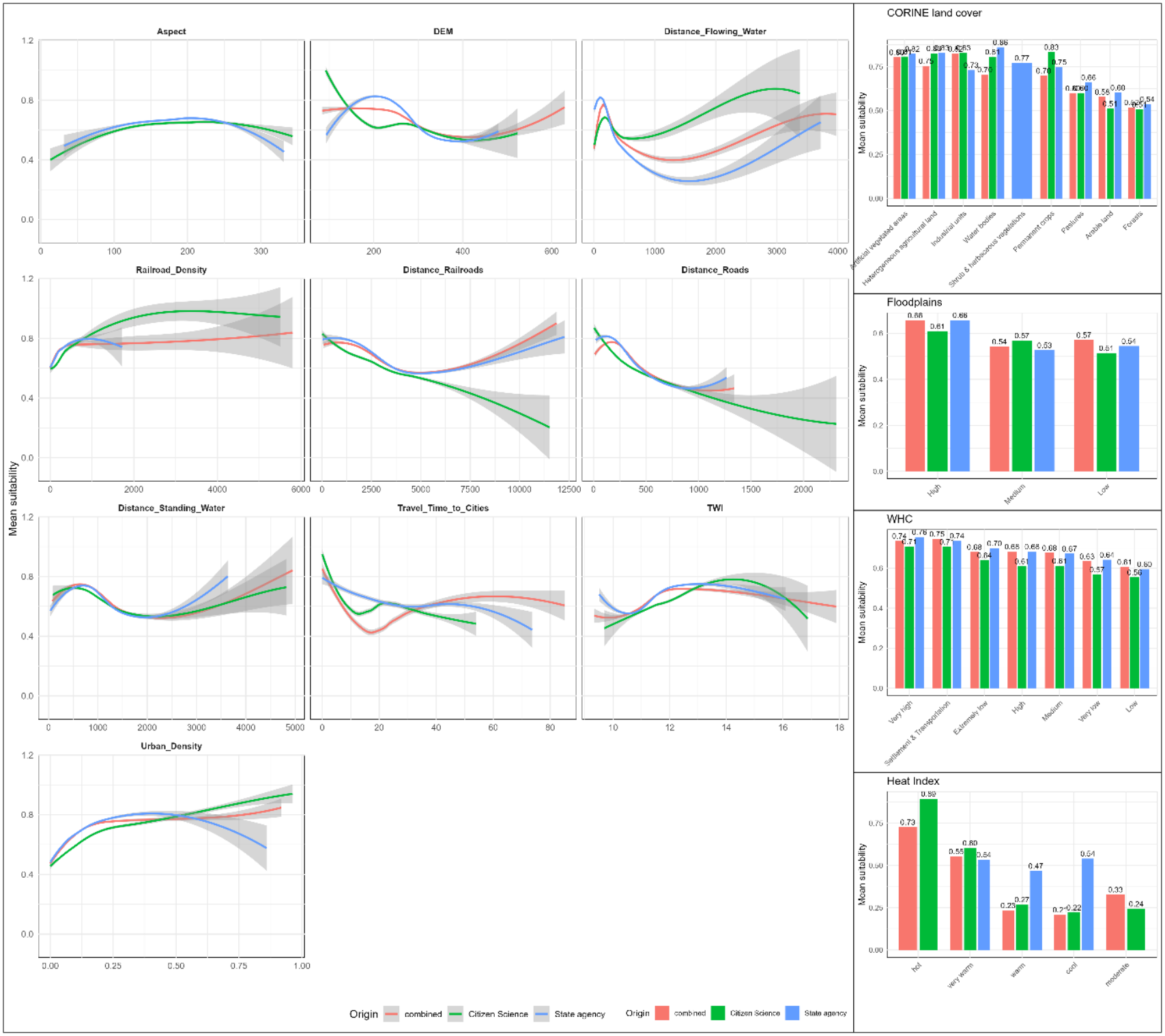

**Appendix 5.2:** Response plots for *Impatiens glandulifera*

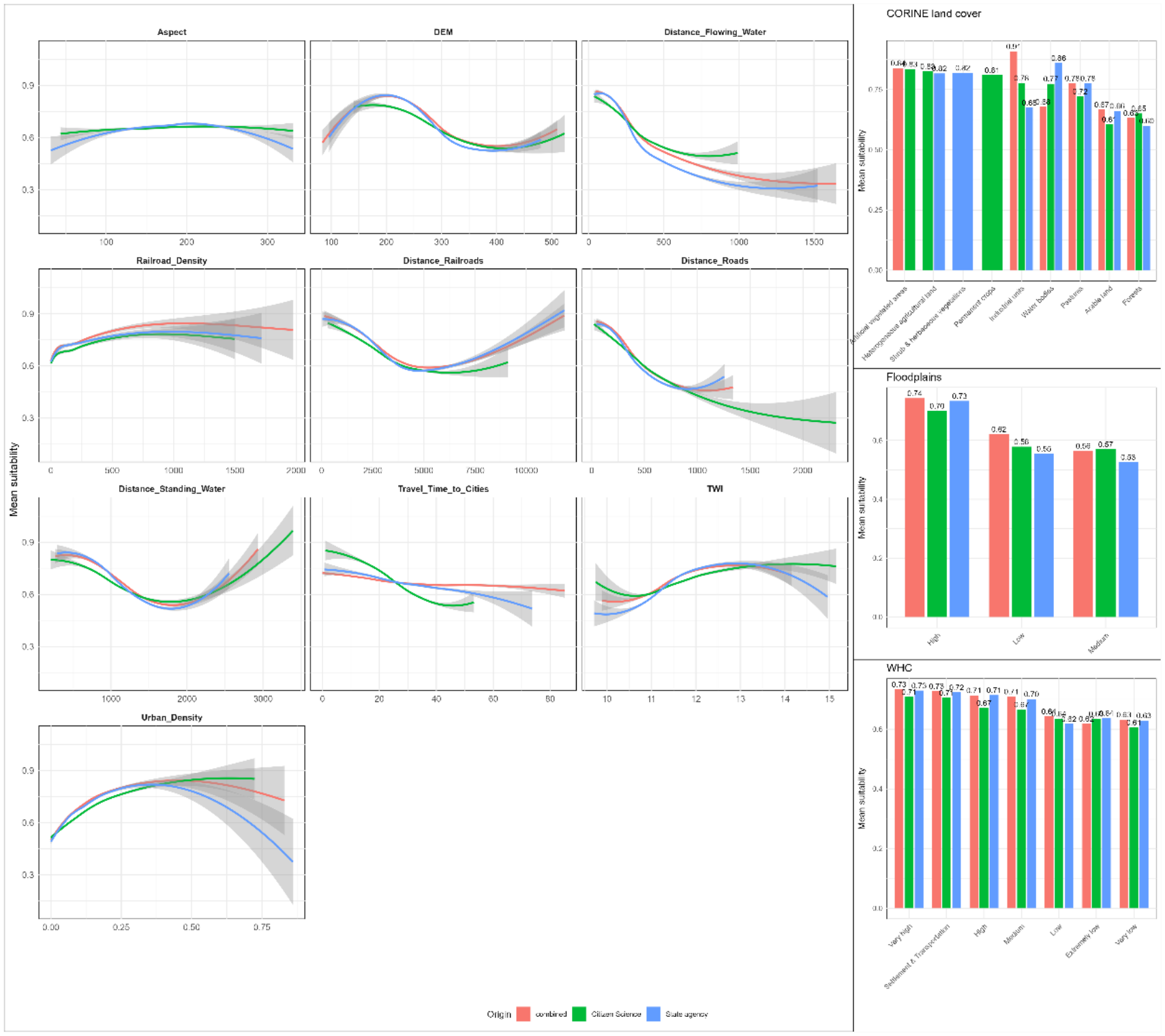

**Appendix 5.3:** Response plots for *Heracleum mantegazzianum*

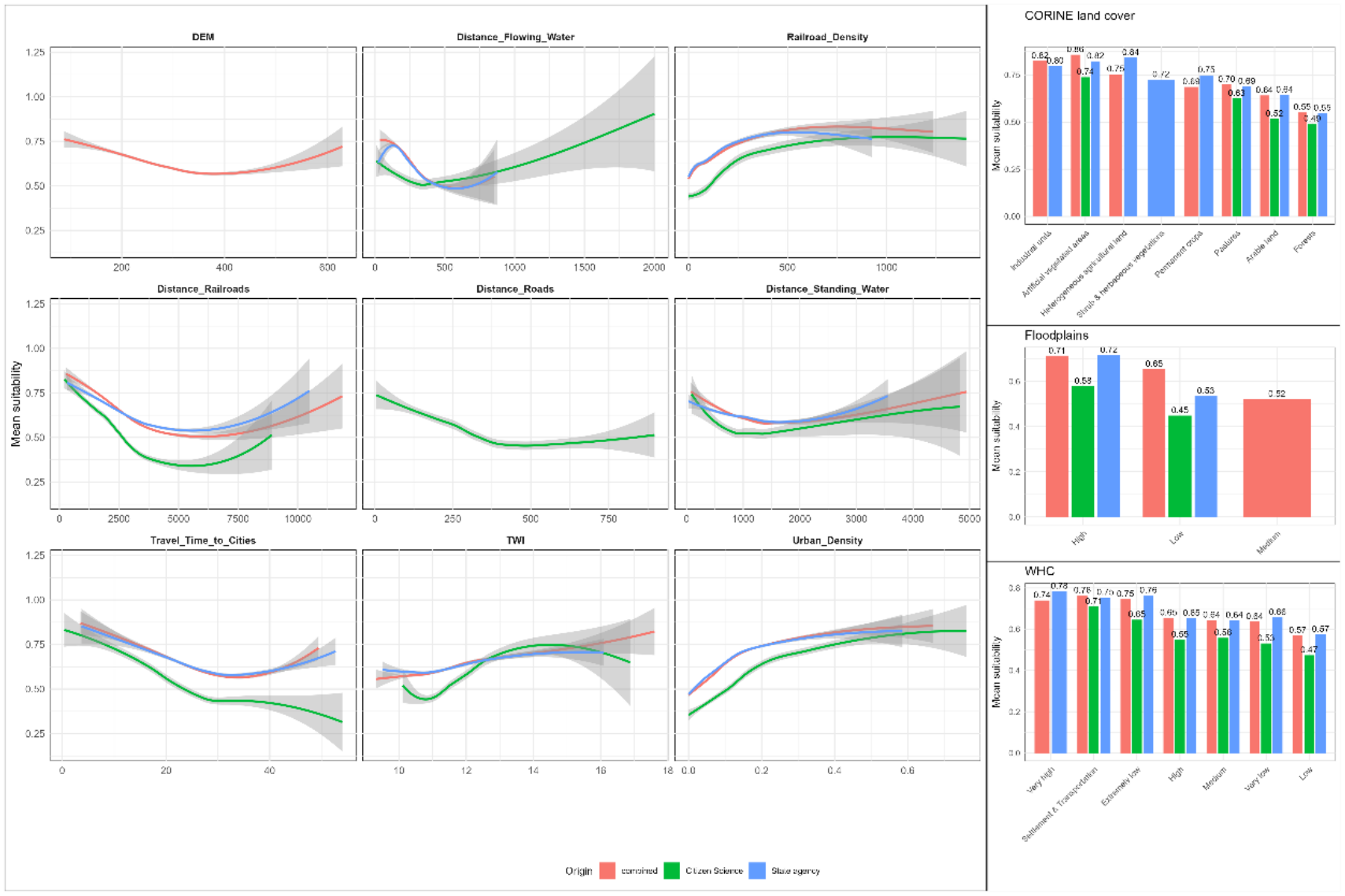

**Appendix 5.4:** Response plots for *Ailanthus altissima*

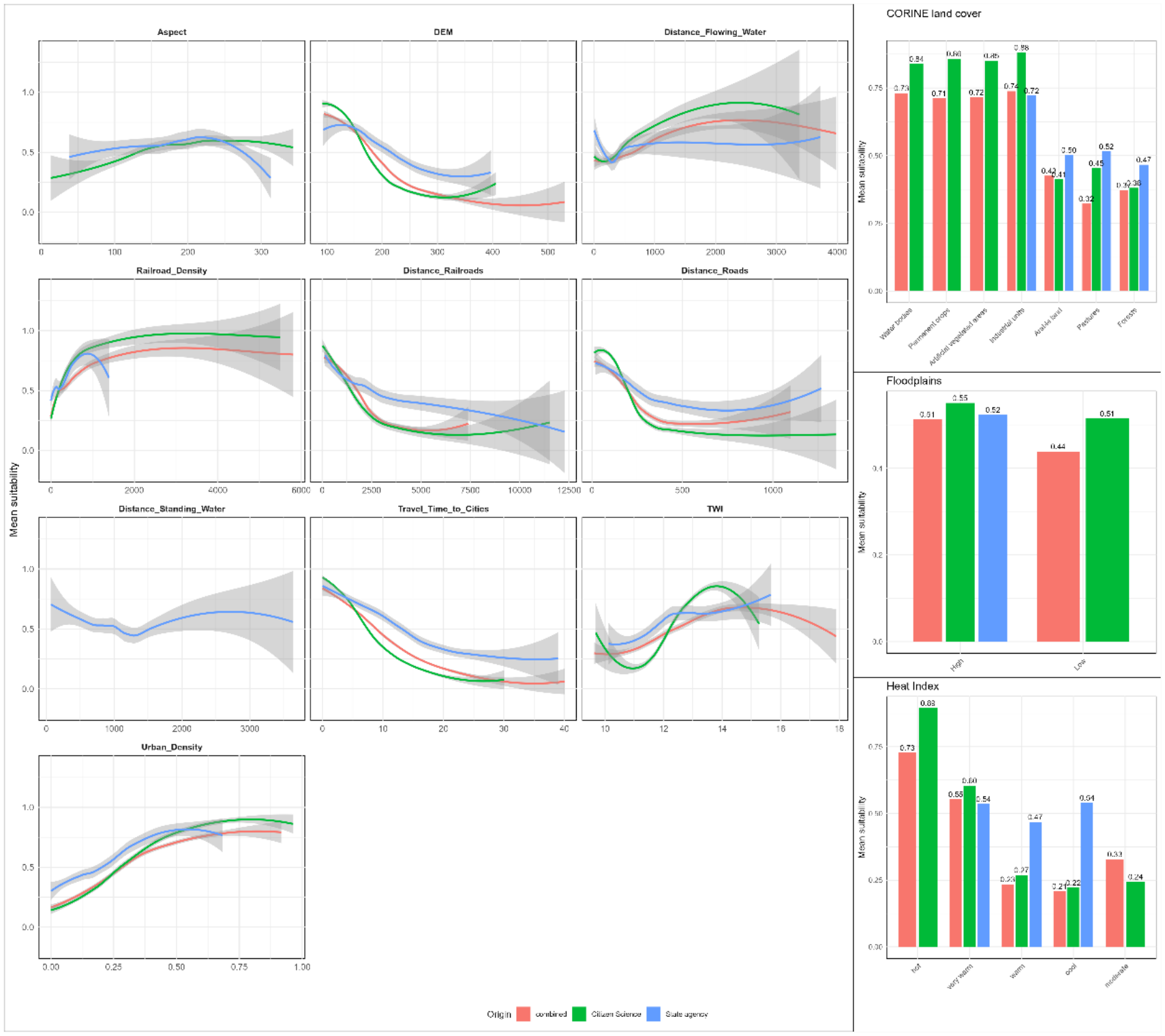

**Appendix 6:** Coefficients of the multiple linear regressions of DI on all predictors per species, averaged over data sources and iterations.

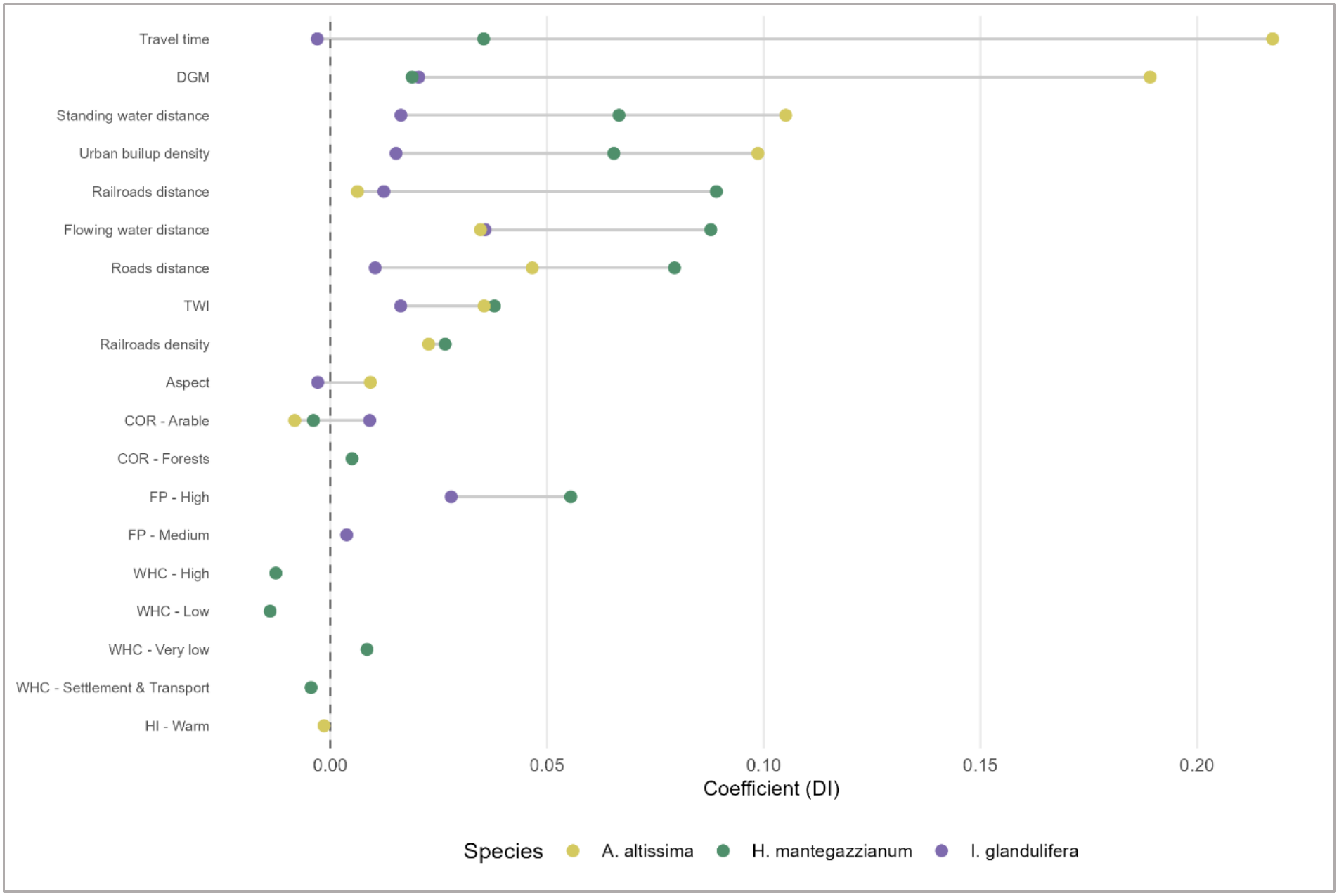

## References

1. Ahmed, N., Atzberger, C., Zewdie, W., 2021. Species Distribution Modelling performance and its implication for Sentinel-2-based prediction of invasive Prosopis juliflora in lower Awash River basin, Ethiopia. Ecol Process 10, 18. 10.1186/s13717-021-00285-6

2. Alexander, J.M., Lembrechts, J.J., Cavieres, L.A., Daehler, C., Haider, S., Kueffer, C., Liu, G., McDougall, K., Milbau, A., Pauchard, A., Rew, L.J., Seipel, T., 2016. Plant invasions into mountains and alpine ecosystems: current status and future challenges. Alp Botany 126, 89–103. 10.1007/s00035-016-0172-8

3. Allouche, O., Tsoar, A., Kadmon, R., 2006. Assessing the accuracy of species distribution models: prevalence, kappa and the true skill statistic (TSS). Journal of Applied Ecology 43, 1223–1232. 10.1111/j.1365-2664.2006.01214.x

4. Anibaba, Ǫ.A., Dyderski, M.K., Jagodziński, A.M., 2022. Predicted range shifts of invasive giant hogweed (Heracleum mantegazzianum) in Europe. Science of The Total Environment 825, 154053. 10.1016/j.scitotenv.2022.154053

5. Barbet-Massin, M., Jiguet, F., Albert, C.H., Thuiller, W., 2012. Selecting pseudo-absences for species distribution models: how, where and how many? Methods Ecol Evol 3, 327–338. 10.1111/j.2041-210X.2011.00172.x

6. Barbet-Massin, M., Rome, Ǫ., Villemant, C., Courchamp, F., 2018. Can species distribution models really predict the expansion of invasive species? PLoS ONE 13, e0193085. 10.1371/journal.pone.0193085

7. Beck, J., Böller, M., Erhardt, A., Schwanghart, W., 2014. Spatial bias in the GBIF database and its effect on modeling species’ geographic distributions. Ecological Informatics 19, 10–15. 10.1016/j.ecoinf.2013.11.002

8. Berio Fortini, L., Kaiser, L.R., Daehler, C.C., Jacobi, J.D., Dimson, M., Gillespie, T.W., 2024. Exploring and integrating differences in niche characteristics across regional and global scales to better understand plant invasions in Hawaiʻi. Biol Invasions 26, 1827–1843. 10.1007/s10530-024-03284-8

9. [dataset] BfG (2024): Water Bodies Germany (Water frameworks directive 3rd cycle) [Wasserkörper-DE (Wasserrahmenrichtlinie 3. Zyklus 2022-2027)]. Available online at https://geoportal.bafg.de/smartfinderClient/?lang=de#/datasets/iso/268d52a5-8409-4fb4-94ed-98f0c7b71d8d.

10. [dataset] BKG (2021). Digital elevation model grid spacing 1000m [Digitales Geländemodell Gitterweite 1000 m] (dgm1000). URL: https://www.bkg.bund.de, licensing: “https://www.govdata.de/dl-de/by-2-0

11. [dataset] BKG (2024): Digitales Landschaftsmodell [Digital Landscape Model] 1:250 000 (DLM250). URL: https://www.bkg.bund.de, licensing: “https://www.govdata.de/dl-de/by-2-0

12. Boria, R.A., Olson, L.E., Goodman, S.M., Anderson, R.P., 2014. Spatial filtering to reduce sampling bias can improve the performance of ecological niche models. Ecological Modelling 275, 73–77. 10.1016/j.ecolmodel.2013.12.012

13. Bracho-Estévanez, C.A., Arenas-Castro, S., González-Varo, J.P., González-Moreno, P., 2024. Spatially explicit metrics improve the evaluation of species distribution models facing sampling biases. Ecological Informatics 84, 102916. 10.1016/j.ecoinf.2024.102916

14. Broennimann, O., Fitzpatrick, M.C., Pearman, P.B., Petitpierre, B., Pellissier, L., Yoccoz, N.G., Thuiller, W., Fortin, M., Randin, C., Zimmermann, N.E., Graham, C.H., Guisan, A., 2012. Measuring ecological niche overlap from occurrence and spatial environmental data. Global Ecology and Biogeography 21, 481–497. 10.1111/j.1466-8238.2011.00698.x

15. Brooks, M., Bolker, B., Kristensen, K., Maechler, M., Magnusson, A., McGillycuddy, M., Skaug, H., Nielsen, A., Berg, C., Bentham, K. van, Sadat, N., Lüdecke, D., Lenth, R., O’Brien, J., Geyer, C.J., Jagan, M., Wiernik, B., Stouffer, D.B., Agronah, M., Akdur, H.T.K., Bové, D.S., Krieger, N., Williams, C., Bogomolovas, J., Purwar, D., 2026. glmmTMB: Generalized Linear Mixed Models using Template Model Builder.

16. [dataset] Brun, P., Zimmermann, N.E., Hari, C., Pellissier, L., Karger, D.N., 2022. CHELSA-BIOCLIM+ A novel set of global climate-related predictors at kilometre-resolution. 10.16904/ENVIDAT.332

17. Cazzolla Gatti, R., Zannini, P., Piovesan, G., Alessi, N., Basset, A., Beierkuhnlein, C., Di Musciano, M., Field, R., Halley, J.M., Hoffmann, S., Iaria, J., Kallimanis, A., Lövei, G.L., Morera, A., Provenzale, A., Rocchini, D., Vetaas, O.R., Chiarucci, A., 2023. Correction to: Analysing the distribution of strictly protected areas toward the EU2030 target. Biodivers Conserv 32, 3175–3177. 10.1007/s10531-023-02683-y

18. César de Sá, N., Marchante, H., Marchante, E., Cabral, J.A., Honrado, J.P., Vicente, J.R., 2019. Can citizen science data guide the surveillance of invasive plants? A model-based test with Acacia trees in Portugal. Biol Invasions 21, 2127–2141. 10.1007/s10530-019-01962-6

19. Cox, T.E., Philippoff, J., Baumgartner, E., Smith, C.M., 2012. Expert variability provides perspective on the strengths and weaknesses of citizen-driven intertidal monitoring program. Ecological Applications 22, 1201–1212. 10.1890/11-1614.1

20. Crall, A.W., Jarnevich, C.S., Young, N.E., Panke, B.J., Renz, M., Stohlgren, T.J., 2015. Citizen science contributes to our knowledge of invasive plant species distributions. Biol Invasions 17, 2415–2427. 10.1007/s10530-015-0885-4

21. Čuda, J., Skálová, H., Pyšek, P., 2020. Spread of *Impatiens glandulifera* from riparian habitats to forests and its associated impacts: insights from a new invasion. Weed Research 60, 8–15. 10.1111/wre.12400

22. Di Cecco, G.J., Barve, V., Belitz, M.W., Stucky, B.J., Guralnick, R.P., Hurlbert, A.H., 2021. Observing the Observers: How Participants Contribute Data to iNaturalist and Implications for Biodiversity Science. BioScience 71, 1179–1188. 10.1093/biosci/biab093

23. Di Febbraro, M., Bosso, L., Fasola, M., Santicchia, F., Aloise, G., Lioy, S., Tricarico, E., Ruggieri, L., Bovero, S., Mori, E., Bertolino, S., 2023. Different facets of the same niche: Integrating citizen science and scientific survey data to predict biological invasion risk under multiple global change drivers. Global Change Biology 29, 5509–5523. 10.1111/gcb.16901

24. Dimson, M., Berio Fortini, L., Tingley, M.W., Gillespie, T.W., 2023. Citizen science can complement professional invasive plant surveys and improve estimates of suitable habitat. Diversity and Distributions 29, 1141–1156. 10.1111/ddi.13749

25. Dormann, C.F., Elith, J., Bacher, S., Buchmann, C., Carl, G., Carré, G., Marquéz, J.R.G., Gruber, B., Lafourcade, B., Leitão, P.J., Münkemüller, T., McClean, C., Osborne, P.E., Reineking, B., Schröder, B., Skidmore, A.K., Zurell, D., Lautenbach, S., 2013. Collinearity: a review of methods to deal with it and a simulation study evaluating their performance. Ecography 36, 27–46. 10.1111/j.1600-0587.2012.07348.x

26. EASIN (2026): European Commission - Joint Research Centre - European Alien Species Information Network (EASIN). Available online at https://easin.jrc.ec.europa.eu/.

27. [dataset] EEA (2019): CORINE Land Cover 2018 (vector), Europe, 6-yearly. version 2020_20u1, May 2020.

28. Feldman, M.J., Imbeau, L., Marchand, P., Mazerolle, M.J., Darveau, M., Fenton, N.J., 2021. Trends and gaps in the use of citizen science derived data as input for species distribution models: A quantitative review. PLoS ONE 16, e0234587. 10.1371/journal.pone.0234587

29. Fourcade, Y., Besnard, A.G., Secondi, J., 2018. Paintings predict the distribution of species, or the challenge of selecting environmental predictors and evaluation statistics. Global Ecol Biogeogr 27, 245–256. 10.1111/geb.12684

30. Gallien, L., Douzet, R., Pratte, S., Zimmermann, N.E., Thuiller, W., 2012. Invasive species distribution models – how violating the equilibrium assumption can create new insights. Global Ecology and Biogeography 21, 1126–1136. 10.1111/j.1466-8238.2012.00768.x

31. [dataset] GBIF (2025a): Occurrence Download - global. 10.15468/dl.ns4htj.

32. [dataset] GBIF (2025b): Occurrence Download - regional. 10.15468/dl.865r6h.

33. Geldmann, J., Heilmann-Clausen, J., Holm, T.E., Levinsky, I., Markussen, B., Olsen, K., Rahbek, C., Tøttrup, A.P., 2016. What determines spatial bias in citizen science? Exploring four recording schemes with different proficiency requirements. Diversity and Distributions 22, 1139–1149. 10.1111/ddi.12477

34. Hartig, F., Lohse, L., 2022. DHARMa: Residual Diagnostics for Hierarchical (Multi-Level / Mixed) Regression Models.

35. Hernandez, P.A., Graham, C.H., Master, L.L., Albert, D.L., 2006. The effect of sample size and species characteristics on performance of different species distribution modeling methods. Ecography 29, 773–785. 10.1111/j.0906-7590.2006.04700.x

36. [dataset] HLNUG. (2024): Species distribution maps [Verbreitungskarten]. Available online at https://www.hlnug.de/themen/naturschutz/tiere-und-pflanzen/arten-melden/invasive-arten/pflanzen/landpflanzen.

37. [dataset] HLNUG (2025a): Water Bodies [Gewässernetz] 1:25.000. Available online at https://www.hlnug.de/themen/geografische-informationssysteme/geodienste/wasser.

38. [dataset] HLNUG (2025b): Usable field capacity in the 1st meter [Nutzbare Feldkapazität des Bodens im 1. Meter zu den Bodenflächendaten] 1:50.000. Available online at https://www.hlnug.de/themen/geografische-informationssysteme/geodienste/boden.

39. [dataset] HLNUG (2025c): Flooding areas [Überflutungsflächen] HǪ10, HǪ100, HǪextrem (HWRMP). Available online at https://www.hlnug.de/themen/geografische-informationssysteme/geodienste/wasser.

40. [dataset] HLNUG (2025d): Occurrence data of Ailanthus altissima, Heracleum mantegazzianum and Impatiens glandulifera. HLNUG Nature Conservation Department. Hessian Biodiversity Database (HEBID). URL: www.hlnug.de/themen/naturschutz/naturschutzdaten

41. Hulme, P.E., Bacher, S., Kenis, M., Klotz, S., Kühn, I., Minchin, D., Nentwig, W., Olenin, S., Panov, V., Pergl, J., Pyšek, P., Roques, A., Sol, D., Solarz, W., Vilà, M., 2008. Grasping at the routes of biological invasions: a framework for integrating pathways into policy. Journal of Applied Ecology 45, 403–414. 10.1111/j.1365-2664.2007.01442.x

42. [dataset] HVBG (2025): Official Topographic and Cartographic Information System [Amtliches Topographisch-Kartographisches Informationssystem] (ATKIS®) – Basis-DLM.

43. Iturbide, M., Bedia, J., Herrera, S., Del Hierro, O., Pinto, M., Gutiérrez, J.M., 2015. A framework for species distribution modelling with improved pseudo-absence generation. Ecological Modelling 312, 166–174. 10.1016/j.ecolmodel.2015.05.018

44. Jiménez-Valverde, A., Lobo, J.M., 2007. Threshold criteria for conversion of probability of species presence to either–or presence–absence. Acta Oecologica 31, 361–369. 10.1016/j.actao.2007.02.001

45. Johnston, A., Moran, N., Musgrove, A., Fink, D., Baillie, S.R., 2020. Estimating species distributions from spatially biased citizen science data. Ecological Modelling 422, 108927. 10.1016/j.ecolmodel.2019.108927

46. Joppa, L.N., Pfaff, A., 2009. High and Far: Biases in the Location of Protected Areas. PLoS ONE 4, e8273. 10.1371/journal.pone.0008273

47. Kiełtyk, P., Delimat, A., 2019. Impact of the alien plant Impatiens glandulifera on species diversity of invaded vegetation in the northern foothills of the Tatra Mountains, Central Europe. Plant Ecol 220, 1–12. 10.1007/s11258-018-0898-z

48. Klinger, Y.P., Eckstein, R.L., Kleinebecker, T., 2023. iPhenology: Using open-access citizen science photos to track phenology at continental scale. Methods in Ecology and Evolution 14. 10.1111/2041-210X.14114

49. Lezcano Caceres, H.L., 2010. ECOLOGICAL CHARACTERISTICS AND ECONOMIC IMPACT OF NON NATIVE Ailanthus altissima (MILL.) SWINGLE IN HESSE, GERMANY. Georg-August-University Göttingen. 10.53846/goediss-2373

50. Liu, C., Newell, G., White, M., 2016. On the selection of thresholds for predicting species occurrence with presence-only data. Ecology and Evolution 6, 337–348. 10.1002/ece3.1878

51. Liu, C., Wolter, C., Xian, W., Jeschke, J.M., 2020. Most invasive species largely conserve their climatic niche. Proc. Natl. Acad. Sci. U.S.A. 117, 23643–23651. 10.1073/pnas.2004289117

52. Lobo, J.M., Jiménez-Valverde, A., Real, R., 2008. AUC: a misleading measure of the performance of predictive distribution models. Global Ecology and Biogeography 17, 145–151. 10.1111/j.1466-8238.2007.00358.x

53. Lozano, V., Chapman, D., Brundu, G., 2017. Native and non-native aquatic plants of South America: comparing and integrating GBIF records with literature data. MBI 8, 443–454. 10.3391/mbi.2017.8.3.18

54. Lozano, V., Marzialetti, F., Acosta, A.T.R., Arduini, I., Bacchetta, G., Domina, G., Laface, V.L.A., Lazzeri, V., Montagnani, C., Musarella, C.M., Nicolella, G., Podda, L., Spampinato, G., Tavilla, G., Brundu, G., 2024. Prioritizing management actions for invasive non-native plants through expert-based knowledge and species distribution models. Ecological Indicators 112279. 10.1016/j.ecolind.2024.112279

55. Mair, L., Ruete, A., 2016. Explaining Spatial Variation in the Recording Effort of Citizen Science Data across Multiple Taxa. PLoS ONE 11, e0147796. 10.1371/journal.pone.0147796

56. Martinez, B., Reaser, J.K., Dehgan, A., Zamft, B., Baisch, D., McCormick, C., Giordano, A.J., Aicher, R., Selbe, S., 2020. Technology innovation: advancing capacities for the early detection of and rapid response to invasive species. Biol Invasions 22, 75–100. 10.1007/s10530-019-02146-y

57. Matutini, F., Baudry, J., Pain, G., Sineau, M., Pithon, J., 2021. How citizen science could improve species distribution models and their independent assessment. Ecology and Evolution 11, 3028–3039. 10.1002/ece3.7210

58. Maxwell, S.L., Fuller, R.A., Brooks, T.M., Watson, J.E.M., 2016. Biodiversity: The ravages of guns, nets and bulldozers. Nature 536, 143–145. 10.1038/536143a

59. Mayer, K., Heger, T., Kühn, I., Nehring, S., Gaertner, M., 2023. Germany’s first Action plan on the pathways of invasive alien species to prevent their unintentional introduction and spread. NB 89, 209–227. 10.3897/neobiota.89.106323

60. McGeoch, M.A., Genovesi, P., Bellingham, P.J., Costello, M.J., McGrannachan, C., Sheppard, A., 2016. Prioritizing species, pathways, and sites to achieve conservation targets for biological invasion. Biol Invasions 18, 299–314. 10.1007/s10530-015-1013-1

61. Meyer, H., Pebesma, E., 2021. Predicting into unknown space? Estimating the area of applicability of spatial prediction models. Methods in Ecology and Evolution 12, 1620– 1633. 10.1111/2041-210X.13650

62. Meyer, H., Reudenbach, C., Wöllauer, S., Nauss, T., 2019. Importance of spatial predictor variable selection in machine learning applications – Moving from data reproduction to spatial prediction. Ecological Modelling 411, 108815. 10.1016/j.ecolmodel.2019.108815

63. Motti, R., Zotti, M., Bonanomi, G., Cozzolino, A., Stinca, A., Migliozzi, A., 2021. Climatic and anthropogenic factors affect Ailanthus altissima invasion in a Mediterranean region. Plant Ecol 222, 1347–1359. 10.1007/s11258-021-01183-9

64. Moudrý, V., Bazzichetto, M., Remelgado, R., Devillers, R., Lenoir, J., Mateo, R.G., Lembrechts, J.J., Sillero, N., Lecours, V., Cord, A.F., Barták, V., Balej, P., Rocchini, D., Torresani, M., Arenas-Castro, S., Man, M., Prajzlerová, D., Gdulová, K., Prošek, J., Marchetto, E., Zarzo-Arias, A., Gábor, L., Leroy, F., Martini, M., Malavasi, M., Cazzolla Gatti, R., Wild, J., Šímová, P., 2024. Optimising occurrence data in species distribution models: sample size, positional uncertainty, and sampling bias matter. Ecography 2024, e07294. 10.1111/ecog.07294

65. Pereira, P. F., Valerio, F., & Lourenço, R. (2023). Incorporating functional connectivity into species distribution models improves the prediction of invasiveness of an exotic species not at niche equilibrium. Biological Invasions, 25(11), 3517–3533. 10.1007/s10530-023-03121-4

66. Pérez, G., Vilà, M., Gallardo, B., 2022. Potential impact of four invasive alien plants on the provision of ecosystem services in Europe under present and future climatic scenarios. Ecosystem Services 56, 101459. 10.1016/j.ecoser.2022.101459

67. Phillips, S.J., Anderson, R.P., Schapire, R.E., 2006. Maximum entropy modeling of species geographic distributions. Ecological Modelling 190, 231–259. 10.1016/j.ecolmodel.2005.03.026

68. Pyšek, P., Hulme, P.E., Simberloff, D., Bacher, S., Blackburn, T.M., Carlton, J.T., Dawson, W., Essl, F., Foxcroft, L.C., Genovesi, P., Jeschke, J.M., Kühn, I., Liebhold, A.M., Mandrak, N.E., Meyerson, L.A., Pauchard, A., Pergl, J., Roy, H.E., Seebens, H., Kleunen, M., Vilà, M., Wingfield, M.J., Richardson, D.M., 2020. Scientists’ warning on invasive alien species. Biol Rev 95, 1511–1534. 10.1111/brv.12627

69. Roberts, D.R., Bahn, V., Ciuti, S., Boyce, M.S., Elith, J., Guillera-Arroita, G., Hauenstein, S., Lahoz-Monfort, J.J., Schröder, B., Thuiller, W., Warton, D.I., Wintle, B.A., Hartig, F., Dormann, C.F., 2017. Cross-validation strategies for data with temporal, spatial, hierarchical, or phylogenetic structure. Ecography 40, 913–929. 10.1111/ecog.02881

70. Robinson, O.J., Ruiz-Gutierrez, Viviana., Reynolds, M.D., Golet, G.H., Strimas-Mackey, M., Fink, D., 2020. Integrating citizen science data with expert surveys increases accuracy and spatial extent of species distribution models. Diversity and Distributions 26, 976–986. 10.1111/ddi.13068

71. Roy, H.E., Pauchard, A., Stoett, P., Renard Truong, T., Bacher, S., Galil, B.S., Hulme, P.E., Ikeda, T., Sankaran, K., McGeoch, M.A., Meyerson, L.A., Nuñez, M.A., Ordonez, A., Rahlao, S.J., Schwindt, E., Seebens, H., Sheppard, A.W., Vandvik, V., Genovesi, P., Wilson, J.R., 2024. IPBES Invasive Alien Species Assessment: Summary for Policymakers. Zenodo. 10.5281/ZENODO.11254974

72. Roy, H.E., Peyton, J., Aldridge, D.C., Bantock, T., Blackburn, T.M., Britton, R., Clark, P., Cook, E., Dehnen-Schmutz, K., Dines, T., Dobson, M., Edwards, F., Harrower, C., Harvey, M.C., Minchin, D., Noble, D.G., Parrott, D., Pocock, M.J.O., Preston, C.D., Roy, S., Salisbury, A., Schönrogge, K., Sewell, J., Shaw, R.H., Stebbing, P., Stewart, A.J.A., Walker, K.J., 2014. Horizon scanning for invasive alien species with the potential to threaten biodiversity in Great Britain. Global Change Biology 20, 3859–3871. 10.1111/gcb.12603

73. Schoener, T.W., 1970. Nonsynchronous Spatial Overlap of Lizards in Patchy Habitats. Ecology 51, 408–418. 10.2307/1935376

74. Seebens, H., Blackburn, T.M., Dyer, E.E., Genovesi, P., Hulme, P.E., Jeschke, J.M., Pagad, S., Pyšek, P., Winter, M., Arianoutsou, M., Bacher, S., Blasius, B., Brundu, G., Capinha, C., Celesti-Grapow, L., Dawson, W., Dullinger, S., Fuentes, N., Jäger, H., Kartesz, J., Kenis, M., Kreft, H., Kühn, I., Lenzner, B., Liebhold, A., Mosena, A., Moser, D., Nishino, M., Pearman, D., Pergl, J., Rabitsch, W., Rojas-Sandoval, J., Roques, A., Rorke, S., Rossinelli, S., Roy, H.E., Scalera, R., Schindler, S., Štajerová, K., Tokarska-Guzik, B., van Kleunen, M., Walker, K., Weigelt, P., Yamanaka, T., Essl, F., 2017. No saturation in the accumulation of alien species worldwide. Nature Communications 8, 14435. 10.1038/ncomms14435

75. Sepulveda, A.J., Dumoulin, C.E., Blanchette, D.L., McPhedran, J., Holme, C., Whalen, N., Hunter, M.E., Merkes, C.M., Richter, C.A., Neilson, M.E., Daniel, W.M., Jones, D.N., Smith, D.R., 2023. When are environmental DNA early detections of invasive species actionable? Journal of Environmental Management 343, 118216. 10.1016/j.jenvman.2023.118216

76. Sladonja, B., Sušek, M., Guillermic, J., 2015. Review on Invasive Tree of Heaven (Ailanthus altissima (Mill.) Swingle) Conflicting Values: Assessment of Its Ecosystem Services and Potential Biological Threat. Environmental Management 56, 1009–1034. 10.1007/s00267-015-0546-5

77. Soultan, A., Safi, K., 2017. The interplay of various sources of noise on reliability of species distribution models hinges on ecological specialisation. PLoS ONE 12, e0187906. 10.1371/journal.pone.0187906

78. Tessarolo, G., Lobo, J.M., Rangel, T.F., Hortal, J., 2021. High uncertainty in the effects of data characteristics on the performance of species distribution models. Ecological Indicators 121, 107147. 10.1016/j.ecolind.2020.107147

79. Thiele, J., Schuckert, U., Otte, A., 2008. Cultural landscapes of Germany are patch-corridor-matrix mosaics for an invasive megaforb. Landscape Ecology 23, 453–465. 10.1007/s10980-008-9202-2

80. Václavík, T., Meentemeyer, R.K., 2012. Equilibrium or not? Modelling potential distribution of invasive species in different stages of invasion. Diversity and Distributions 18, 73–83. 10.1111/j.1472-4642.2011.00854.x

81. Vorstenbosch, T., Essl, F., Lenzner, B., 2020. An uphill battle? The elevational distribution of alien plant species along rivers and roads in the Austrian Alps. NB 63, 1–24. 10.3897/neobiota.63.55096

82. [dataset] Wan, Z.; Hook, S.; Hulley, G. (2015): MOD11A2 MODIS/Terra Land Surface Temperature/Emissivity 8-Day L3 Global 1km SIN Grid V006. NASA EOSDIS Land Processes DAAC. Data Processing: HLNUG - Remote Sensing Competence Center. Downloaded from HLNUG https://umweltdaten.hessen.de/klima/geodaten/

83. Warren, D.L., Glor, R.E., Turelli, M., 2008. ENVIRONMENTAL NICHE EǪUIVALENCY VERSUS CONSERVATISM: ǪUANTITATIVE APPROACHES TO NICHE EVOLUTION. Evolution 62, 2868–2883. 10.1111/j.1558-5646.2008.00482.x

84. Weiss, D.J., Nelson, A., Gibson, H.S., Temperley, W., Peedell, S., Lieber, A., Hancher, M., Poyart, E., Belchior, S., Fullman, N., Mappin, B., Dalrymple, U., Rozier, J., Lucas, T.C.D., Howes, R.E., Tusting, L.S., Kang, S.Y., Cameron, E., Bisanzio, D., Battle, K.E., Bhatt, S., Gething, P.W., 2018. A global map of travel time to cities to assess inequalities in accessibility in 2015. Nature 553, 333–336. 10.1038/nature25181

85. Zając, A., Tokarska-Guzik, B., Zając, M., 2011. The role of rivers and streams in the migration of alien plants into the Polish Carpathians. Biodiversity: Research and Conservation 23, 43–56. 10.2478/v10119-011-0012-z

86. Zizka, A., Silvestro, D., Andermann, T., Azevedo, J., Duarte Ritter, C., Edler, D., Farooq, H., Herdean, A., Ariza, M., Scharn, R., Svantesson, S., Wengström, N., Zizka, V., Antonelli, A., 2019. COORDINATECLEANER : Standardized cleaning of occurrence records from biological collection databases. Methods Ecol Evol 10, 744–751. 10.1111/2041-210X.13152

## References

87. Čuda, J., Skálová, H., Pyšek, P., 2020. Spread of Impatiens glandulifera from riparian habitats to forests and its associated impacts: insights from a new invasion. Weed Research 60, 8–15. 10.1111/wre.12400

88. HLNUG, 2018. Beobachteter Klimawandel. Klimawandel in Hessen. Hessisches Landesamt für Naturschutz, Umwelt und Geologie, Wiesbaden. https://www.hlnug.de/fileadmin/dokumente/klima/beobachteter_klimawandel.pdf

89. HMLU, 2024. Wir machen den Wald klimastabil. Hessisches Ministerium für Landwirtschaft und Umwelt, Weinbau, Forsten, Jagd und Heimat. https://landwirtschaft.hessen.de/wald (accessed 08.10.2026).

90. Huebener, H., Gelhardt, U., Lang, J., 2022. Improved representativeness of simulated climate using natural units and monthly resolution. Frontiers in Climate 4, 991082. 10.3389/fclim.2022.991082

91. Jauker, F., Diekötter, T., Schwarzbach, F., Wolters, V., 2009. Pollinator dispersal in an agricultural matrix: opposing responses of wild bees and hoverflies to landscape structure and distance from main habitat. Landscape Ecology 24, 547–555. 10.1007/s10980-009-9331-2

92. [dataset] NASA JPL, 2020. NASADEM Merged DEM Global 1 arc second V001. NASA EOSDIS Land Processes DAAC. 10.5067/MEaSUREs/NASADEM/NASADEM_HGT.001

93. Reger, B., Mattern, T., Otte, A., Waldhardt, R., 2009. Assessing the spatial distribution of grassland age in a marginal European landscape. Journal of Environmental Management 90, 2900–2909. 10.1016/j.jenvman.2007.10.015

94. Sladonja, B., Sušek, M., Guillermic, J., 2015. Review on Invasive Tree of Heaven (Ailanthus altissima (Mill.) Swingle) Conflicting Values: Assessment of Its Ecosystem Services and Potential Biological Threat. Environmental Management 56, 1009–1034. 10.1007/s00267-015-0546-5

95. Statistisches Bundesamt, 2024. Fläche und Bevölkerung nach Ländern. https://www.statistikportal.de/de/bevoelkerung/flaeche-und-bevoelkerung (accessed 08.10.2026).

96. Thiele, J., Schuckert, U., Otte, A., 2008. Cultural landscapes of Germany are patch-corridor-matrix mosaics for an invasive megaforb. Landscape Ecology 23, 453–465. 10.1007/s10980-008-9202-2

## References

97. [dataset] Brun, P., Zimmermann, N.E., Hari, C., Pellissier, L., Karger, D.N., 2022. CHELSA-BIOCLIM+ A novel set of global climate-related predictors at kilometre-resolution. 10.16904/ENVIDAT.332

98. [dataset] GBIF (2025a): Occurrence Download - global. 10.15468/dl.ns4htj.

99. [dataset] GBIF (2025b): Occurrence Download - local. 10.15468/dl.865r6h.

100. Zizka, A., Silvestro, D., Andermann, T., Azevedo, J., Duarte Ritter, C., Edler, D., Farooq, H., Herdean, A., Ariza, M., Scharn, R., Svantesson, S., Wengström, N., Zizka, V., Antonelli, A., 2019. COORDINATECLEANER : Standardized cleaning of occurrence records from biological collection databases. Methods Ecol Evol 10, 744–751. 10.1111/2041-210X.13152

## References

101. [dataset] Brun, P., Zimmermann, N.E., Hari, C., Pellissier, L., Karger, D.N., 2022. CHELSA-BIOCLIM+ A novel set of global climate-related predictors at kilometre-resolution. 10.16904/ENVIDAT.332

102. [dataset] EEA (2019): CORINE Land Cover 2018 (vector), Europe, 6-yearly. Version 2020_20u1, May 2020.

103. [dataset] HLNUG (2025b): Usable field capacity in the 1st meter [Nutzbare Feldkapazität des Bodens im 1. Meter zu den Bodenflächendaten] 1:50.000. Available online at https://www.hlnug.de/themen/geografische-informationssysteme/geodienste/boden.

104. [dataset] Wan, Z.; Hook, S.; Hulley, G. (2015): MOD11A2 MODIS/Terra Land Surface Temperature/Emissivity 8-Day L3 Global 1km SIN Grid V006. NASA EOSDIS Land Processes DAAC. Data Processing: HLNUG - Remote Sensing Competence Center. Downloaded from HLNUG https://umweltdaten.hessen.de/klima/geodaten/

